# Unlimited cooperativity of *Betatectivirus* SSB, a novel DNA binding protein related to an atypical group of SSBs from protein-primed replicating bacterial viruses

**DOI:** 10.1101/2021.03.25.437074

**Authors:** A. Lechuga, D. Kazlauskas, M. Salas, M. Redrejo-Rodríguez

## Abstract

Bam35 and related betatectiviruses are tail-less bacteriophages that prey on members of the *Bacillus cereus* group. These temperate viruses replicate their linear genome by a protein-primed mechanism. In this work, we have identified and characterized the product of the viral ORF2 as a single-stranded DNA binding protein (hereafter B35SSB). B35SSB binds ssDNA with great preference over dsDNA or RNA in a sequence-independent, highly cooperative manner that results in a non-specific stimulation of DNA replication. We have also identified several aromatic and basic residues, involved in base-stacking and electrostatic interactions, respectively, that are required for effective protein-ssDNA interaction.

Although SSBs are essential for DNA replication in all domains of life as well as many viruses, they are very diverse proteins. However, most SSBs share a common structural domain, named OB-fold. Protein-primed viruses could constitute an exception, as no OB-fold DNA binding protein has been reported. Based on databases searches as well as phylogenetic and structural analyses, we showed that B35SSB belongs to a novel and independent group of SSBs. This group contains proteins encoded by protein-primed viral genomes from unrelated viruses, spanning betatectiviruses and Φ29 and close podoviruses, and they share a conserved pattern of secondary structure. Sensitive searches and structural predictions indicate that B35SSB contains a conserved domain resembling a divergent OB-fold, which would constitute the first occurrence of an OB-fold-like domain in a protein-primed genome.

**Highlights:**

- Bam35 ORF 2 product encodes a viral single-stranded DNA binding protein (B35SSB).
- B35SSB binds ssDNA in a highly cooperative manner but with no sequence specificity.
- B35SSB-ssDNA binding is mediated by base-stacking and ionic interactions.
- Bam35 and Φ29-related SSBs form a novel group of SSBs from protein-primed viruses.
- The B35-Φ29 SSBs group shares a highly divergent OB-fold-like domain.

## Introduction

The *Bacillus* virus Bam35 is the model virus of the genus *Betatectivirus*, a group of temperate *Tectiviridae* members infecting Gram-positive animal and human pathogens from the *Bacillus cereus* group [1,2]. Besides preying on pathogens of economic and global health relevance, this group of bacteriophages has also raised interest in the last years after the suggestion of a possible evolutionary link between betatectiviruses and the origin of several groups of large DNA viruses [3,4].

The genome of Bam35 is replicated by a protein-priming DNA replication process [5], a widespread mechanism for the initiation of genome replication in a number of linear genomes of viruses and linear plasmids. By this mechanism, a specific amino acid of the so-called terminal protein (TP) primes the replication providing a hydroxyl group for the incorporation of the first nucleotide by the viral DNA polymerase and thus it becomes covalently linked to the 5’ genome ends. Among the characterized models of protein-primed DNA replication, the *Bacillus* virus Φ29 from the *Podoviridae* family has been extensively characterized [6,7]. Notwithstanding clear mechanistic differences, Bam35 genome replication can be carried out *in vitro* with only two proteins, the DNA polymerase (B35DNAP) and the TP (B35TP), as in the case of genome replication of Φ29 or PRD1, a well-characterized lytic tectivirus from the genus *Alphatectivirus* infecting Gram-negative hosts [8]. However, although not identified in Bam35, a number of accessory DNA binding proteins, such as single-stranded DNA binding proteins (SSBs), increase the efficiency of genome replication *in vitro* and are essential *in vivo* in the case of Φ29 and other systems [9–11].

SSBs are ubiquitous factors that protect single-stranded DNA (ssDNA) intermediates required for genetic information metabolism. These proteins not only protect against nucleases attack, breakage and chemical mutagens, but also can play active roles in preventing secondary structures in the ssDNA, recruiting enzymes and stimulating DNA replication by enhancing the processivity, rate and fidelity of DNA synthesis [12–15]. Warranted by all those features, SSBs have also proved to be suitable for diverse molecular biology and analytical applications in biotechnology [16–18].

Although SSBs span a wide diversity of protein groups with little sequence similarity, the vast majority of them share a common structural domain, called oligonucleotide/oligosaccharide-binding fold (OB-fold). This domain consists of a five-stranded β-sheet coiled to form a β-barrel capped by an α-helix [19–21]. There are very few exceptions of SSB structures with no clear similarity to an OB-fold, such as those in adenovirus [22], the hyperthermophilic archaea representing clade Thermoproteales [23] and Drc protein from the N4-related bacterial viruses [24].

Generally, OB-fold domains interact with ssDNA by base stacking with aromatic residues situated in strands 2 and 3 of the β-barrel. Also, cation-π stacking, hydrophobic and hydrogen-bonding with base and ribose moieties contribute to the SSB-bases binding without sequence specificity. Meanwhile, the phosphate backbone is often exposed to the solvent, although, it can also contribute to ssDNA binding through salt bridges and hydrogen bonds [25].

As each OB-fold unit can bind a very short ssDNA tract, SSBs present different modular organization, either by the presence of more than one OB-fold in the same polypeptide, as in the case of eukaryotic RPA, or by oligomerization of independent OB-fold monomers, often by interaction among their C-terminal tails [26]. In other cases, the N-terminal mediates oligomerization, as in the case of GA-1 viral SSB, which is a hexamer with a highly efficient DNA binding ability, whereas the Φ29SSB is a monomer with lower DNA binding proficiency [27]. Furthermore, SSB-ssDNA interaction may be required to trigger multimerization complex formation. This can result in a cooperative DNA binding mechanism, common for SSBs whose main function is related to DNA replication, such as recA, the SSBs from phage T4 or *E. coli* (EcoSSB), among others [28,29]. However, cooperativity can be also low in some SSBs, as T7 gp2.5 and eukaryotic RPA [30,31]. Further, some SSBs, like EcoSSB, can show either limited or unlimited cooperative binding depending on the salt concentration [32].

In this work, we identified and characterized the product of the Bam35 ORF 2 (hereafter B35SSB) as a novel viral SSB. Biochemical characterization of B35SSB reveals high specificity for ssDNA binding in a greatly cooperative manner, which results in the stimulation of processive DNA replication. Site-directed mutagenesis also allowed us to disclose some aspects of the DNA binding mechanism, similar to that reported for other SSBs with a canonical OB-fold. Further, phylogenetic analyses and consensus structural predictions showed that this protein belongs to a diverse clade of SSBs that contains, besides betatectivirus orthologs, Φ29-related podoviruses and diverse bacterial proteins that might be also coded by uncharacterized or overlooked viral genomes. This group of viral SSBs would constitute a novel clade of SSBs from protein-primed replicating genomes that would contain an OB-fold-like conserved domain.

## RESULTS

### B35SSB is a proficient single-stranded DNA binding protein

Characterization of the protein P2 from the betatectivirus-related *B. cereus* plasmid pBClin15, orthologous to the Bam35 ORF2 product, showed DNA binding capacity [33]. Other works proposed that Bam35 ORF2 encodes an SSB similar to other bacteriophage SSBs based on HHpred searches [34]. To confirm that the product of the Bam35 ORF2 is a viral SSB protein, we analyzed its nucleic acid binding ability by electrophoretic mobility shift assay (EMSA) using a variety of nucleic acid substrates. First, a 50-mer ssDNA and dsDNA oligonucleotides were incubated with increasing amounts of purified protein (hereafter B35SSB). The SSB from *E. coli* (EcoSSB) and the TP from Φ29 were used as positive controls (Figure 1A), as the EcoSSB binds more efficiently ssDNA (lanes 1 and 8) while the TP shows higher affinity for dsDNA (lanes 2 and 9). In presence of B35SSB, the ssDNA substrate showed slower mobility due to the formation of B35SSB-ssDNA stable complexes (lanes 4-6). The lack of intermediate bands suggests a cooperative DNA binding. Moreover, although the EcoSSB monomer has a similar molecular weight to B35SSB (19 kDa and 18.5 kDa, respectively), EcoSSB-ssDNA complexes (lane 2) showed higher mobility than B35SSB-ssDNA complexes (lane 6), which indicates that a higher number of B35SSB molecules are bound to the ssDNA fragment.

**Figure 1.**
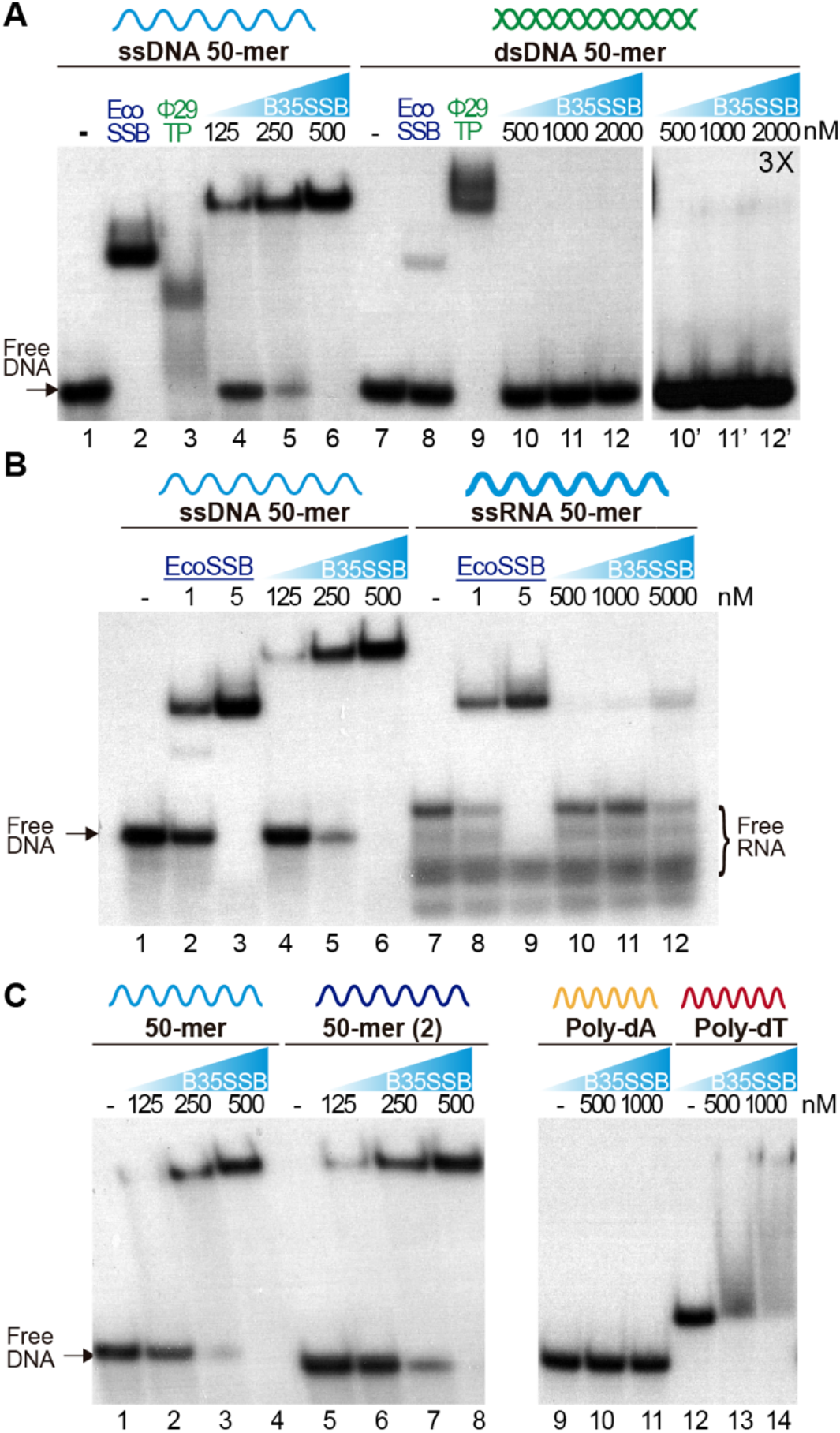
Nucleic acid binding capacity of B35SSB. DNA binding capacity of B35SSB is analyzed by comparison of mobility shift of diverse substrates, ssDNA vs. dsDNA (A), ssDNA vs. ssRNA (B) or different sequence contexts in ssDNA (C). EMSA were performed in presence of the indicated amounts of B35SSB and 2 nM of radiolabeled ssDNA, dsDNA or RNA as indicated. Negative controls, without protein, were loaded in the lanes indicated by a dash. Where indicated, 1μM EcoSSB and 20 nM Φ29 terminal protein (Φ29TP) were used as controls.

Participation of SSBs in DNA replication initiation steps has been related to a binding capacity to RNA-DNA or to duplex dsDNA [10,35]. Nevertheless, dsDNA binding of B35SSB seems insignificant as only a very faint shifted band was observed (lane 12’) at high a concentration of B35SSB (1000:1, B35SSB:dsDNA molar ratio), whereas B35SSB could shift nearly all ssDNA substrate at a 125:1 (B35SSB:ssDNA) molar ratio (lane 5). Recent works have also highlighted the RNA binding capacity of several SSBs from bacteria and viruses, leading to RNA metabolism regulation [24,36]. However, when RNA binding capacity of B35SSB was analyzed, only a single weak band was detected at a very high concentration of B35SSB (Figure 1B, lanes 7-12) indicating an inefficient binding to RNA, as compared with ssDNA of the equivalent sequence (Figure 1B, lanes 1-6). While EcoSSB-ssDNA and EcoSSB-RNA complexes migrate a similar distance, B35SSB-RNA migrated faster than B35SSB-ssDNA complexes. This suggests a different mode of binding of B35SSB to RNA where a smaller number of molecules could bind to the RNA substrate with less affinity than to ssDNA. Therefore, we conclude that B35SSB binds ssDNA proficiently, with a great preference over dsDNA or RNA substrates.

We also analyzed different ssDNA substrates to determine the sequence-specificity in DNA binding of B35SSB. Overall, no differences were observed when two different 50-mer ssDNA substrates were used (Figure 1C, lines 2-4 vs. 6-8). However, when we compared homopolymeric 33-mer substrates, we found that B35SSB can bind poly-dT oligonucleotides (lines 12-14), but its binding capacity is severely impaired on poly-dA homopolymeric ssDNA (lines 9-11). This indicates a preference for pyrimidines over purines, as previously reported for other OB-fold containing SSBs [12], suggesting a similar nature of interactions involved in B35SSB binding mechanism to ssDNA. Altogether, these results confirm the product of Bam35 ORF 2 as a viral SSB, with an intrinsic high affinity for ssDNA and low sequence specificity, but a negligible capacity of binding to other substrates.

### B35SSB stimulates processive DNA replication

To ascertain the role of B35SSB in DNA replication, we next determined its influence during processive DNA replication in a rolling circle replication assay *in vitro*. Thus, we used a singly primed ssDNA of M13 as a template, which can be replicated by Bam35 DNA polymerase (B35DNAP), giving rise to a replication product larger than full-length M13 DNA thanks to its ability to couple processive DNA replication and strand displacement capacity (Figure 2A) [37]. When increasing amounts of SSB were added to the reaction, a stimulatory effect of B35SSB was observed in terms of a higher amount of DNA replication product (Figure 2B, lanes 2-3). However, in the presence of very high concentrations of B35SSB (lanes 4-5), a general decrease of ssDNA product was observed, eventually resulting in the strong reduction of product larger than full-length M13 DNA. Thus, B35SSB stimulates DNA replication by B35DNAP in these conditions, although a fine-tuned protein:DNA ration may be required. This stimulatory effect was also observed in a time-course experiment where B35SSB effect could be detected even at the shortest reaction times of replication (Figure 2C). This could suggest a possible increase in the elongation rate in the presence of B35SSB during DNA replication. Interestingly, rather than specific, the stimulation of its cognate DNA polymerase by B35SSB seems generalized. As such, B35SSB can stimulate also Φ29 DNA polymerase and the B35DNAP can be stimulated by *E. coli* and Φ29 SSBs, although the optimum SSB:DNA ratio might be different (Figure 2D-E).As in the case of Φ29SSB [38], the stimulation of processive DNA replication by B35SSB seems to be facilitated by a DNA helix-destabilizing activity (Figure S1-A), which further supports a role in genome replication. Although this DNA unwinding capacity can be also required at replication initiation steps [39], B35SSB seems irrelevant for protein-primed DNA replication initiation *in vitro* (Figure S1-B-D).Therefore, we conclude that rather than in early steps of protein-primed genome replication, B35SSB would play a role in the processive genome replication.

**Figure 2.**
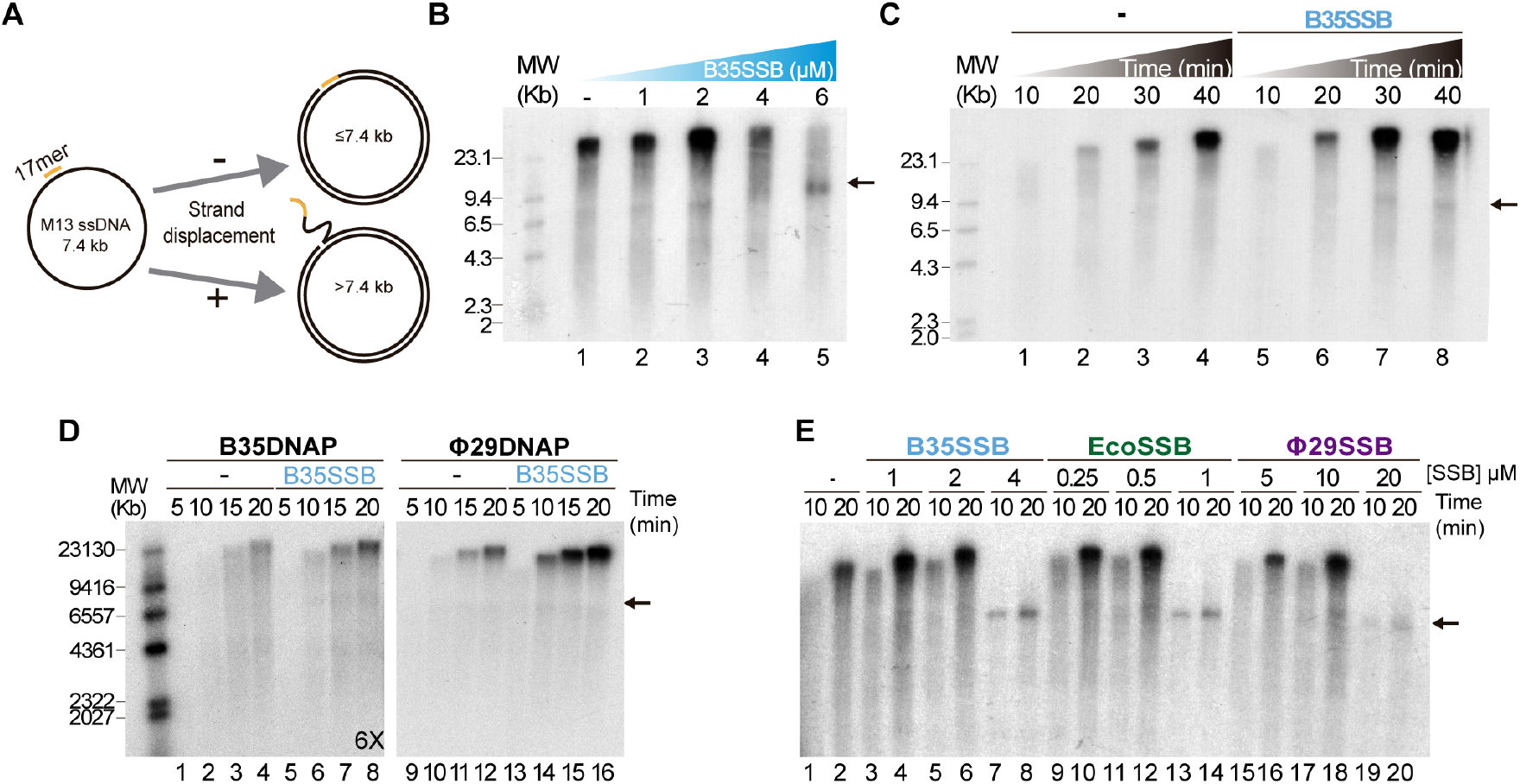
B35SSB stimulation of processive DNA replication. **(A)** Schematic representation of M13 ssDNA rolling-circle replication. In strand-displacement conditions, a product much larger than the full-length M13-DNA (7.4 kb) is obtained. **(B)** Effect of B35SSB concentration in ssDNA M13 replication. **(C)** Time-course analysis of addition of B35SSB. Reactions in absence or presence of 2 μM B35SSB were carried out and stopped at the indicated times. **(D)** Stimulatory capacity of B35SSB (2 μM) on M13 DNA replication performed by Bam35 and Φ29 DNA polymerases. Longer autoradiography exposition time is indicated. **(E)** Effect of different SSBs on B35DNAP processive DNA replication.

Reactions were carried out using primed M13 circular ssDNA as template and the indicated DNA polymerase. After incubation at 37 °C, the length of the synthesized DNA was analyzed by alkaline 0.7% agarose gel electrophoresis alongside a λ DNA (MW) and autoradiography. M13 ssDNA unit length is indicated by a black arrow. See Methods for details.

### B35SSB is a monomer that binds ssDNA in a highly cooperative manner

To analyze the DNA binding mechanism of B35SSB, we first checked the effect of ssDNA length. As shown in Figure 3A, whereas shorter DNA fragments require a higher protein amount to give rise to a DNA shift, producing some smear of unstable protein-DNA complexes in the conditions assayed (lanes 1-7), larger DNA tracts, above 50 nucleotides, are bound stably with lower protein concentration (lanes 8-15). These results indicate that B35SSB binds ssDNA in a highly cooperative way that would be enhanced by the length of the substrate. Moreover, contrary to other SSBs [40,41], B35SSB cooperativity is not affected by the ionic strength, as high salt concentrations impair DNA binding moderately without affecting the pattern of retarded DNA or leading to smeared or intermediate products (Figure 3B).

**Figure 3.**
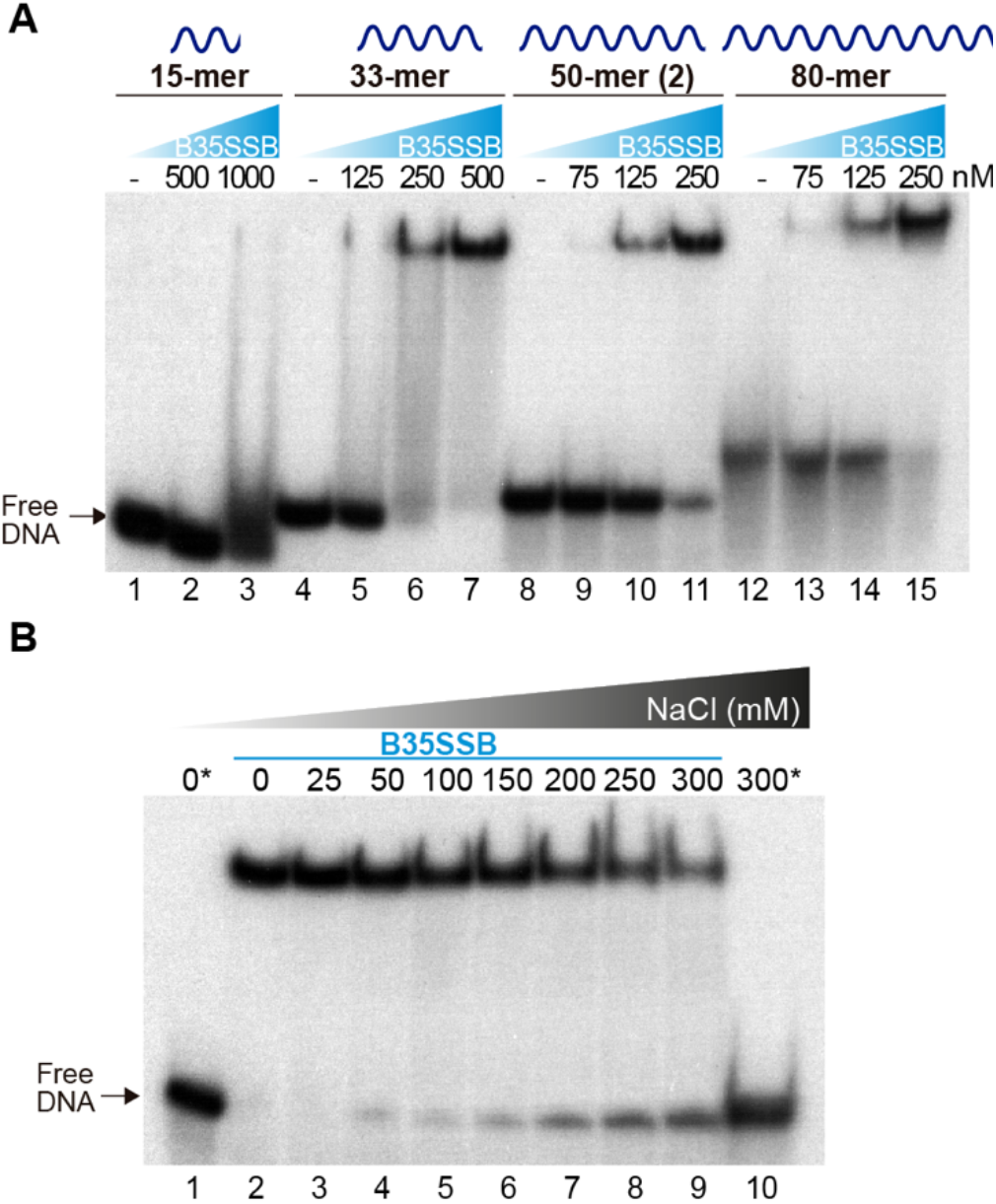
Characterization of cooperative DNA binding by EMSA. (A) Effect of substrate DNA length in ssDNA binding as determined by oligonucleotides of the indicated length. Note that a lower B35SSB concentration range was used for longer ssDNA fragments. (B) DNA binding assay in the presence of increasing concentrations of NaCl. Samples with no protein are indicated with an *.

Cooperative DNA binding may also depend on protein-protein interactions that facilitate the formation of long, stable protein-DNA complexes. We examined the oligomerization state of B35SSB in solution by analytical ultracentrifugation in a linear 15-30% glycerol gradient. The sedimentation peak of B35SSB revealed that it is homogeneously a monomer in solution (Figure 4A). This suggests that oligomerization would occur and be triggered by DNA binding.

**Figure 4.**
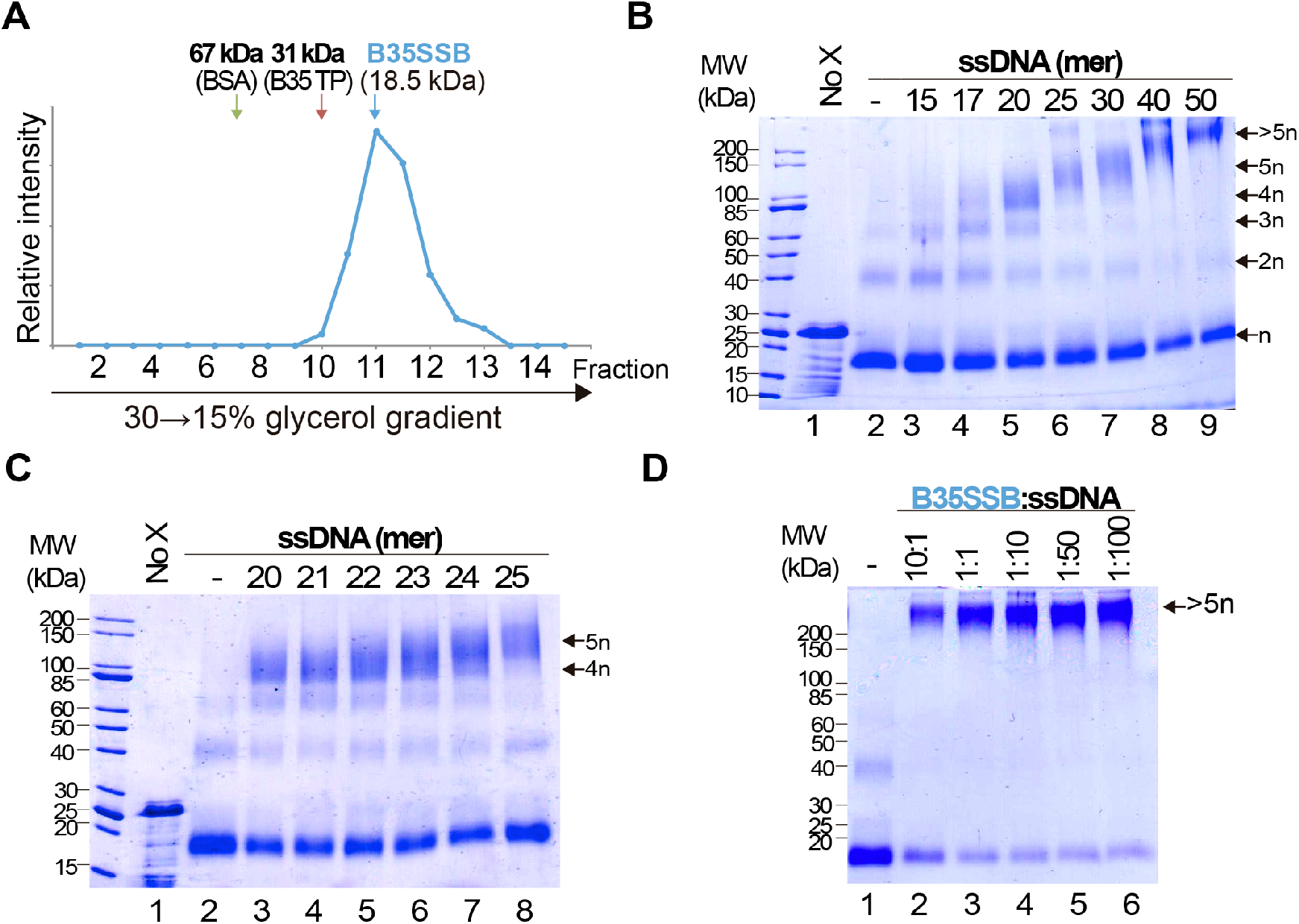
Analysis of B35SSB modularity and cooperativity. **(A)** Determination of the oligomerization state of B35SSB in solution. Recombinant B35SSB was subjected to sedimentation in 15-30% (w/v) glycerol gradients in the presence of molecular weight markers as indicated. The fractions at which the maximal amount of each protein appears are indicated by arrows. **(B)** Cross-linking protein interaction analysis using different ssDNA sizes (Table S1). B35SSB (1 μM, equivalent to 2 μg) was incubated with 165 ng of the indicated oligonucleotides. After glutaraldehyde cross-linking, total protein was precipitated and analyzed by SDS-PAGE. The number of SSB cross-linked species is indicated on the right. **(C)** Determination of the B35SSB minimal binding site by crosslinking assays. **(D)** Effect of B35SSB:ssDNA ratio in ssDNA-SSB binding. Assays contained 1 μg B35SSB and increasing amounts of the 50-mer oligonucleotide to obtain the indicated protein:DNA ratio. See Methods for details. MW, molecular weight; No X and dash (-) stand for no crosslinker and no ssDNA lanes, respectively.

We also performed glutaraldehyde cross-linking assays to disclose the cooperative DNA binding mechanism [42,43]. In the absence of ssDNA, and, in agreement with the previous result, cross-linked B35SSB is still a monomer (Figure 4B, lane 2). Only a minor band corresponding to dimeric species was detected in presence of the cross-linker, likely due to the stabilization of weak or transient protein-protein interactions that disappear in the presence of large ssDNA fragments. On the other hand, in the presence of ssDNA of increasing lengths, diverse oligomeric species can be detected (lanes 3-6). Longer substrates resulted in higher bands that would correspond to oligomers beyond the resolution of the gel. A detailed analysis with DNA fragments from 20 to 25-mer, increasing one nucleotide at a time, allowed us to determine the minimal binding site of B35SSB (Figure 4C). While with 20-23 mer substrates the main oligomeric species corresponded to a tetramer (lanes 3-6), when 24 and 25-mer substrates were added, the tetramer band lost intensity and the band corresponding to the pentameric species was detected (lanes 7 and 8). This indicates that B35SSB has a binding site that can span 4-5 nt. Altogether, these results show that B35SSB is a monomer in solution, and it forms oligomers as it binds to ssDNA cooperatively with a minimal binding site of 4-5 nucleotides per monomer, being cooperativity stimulated when longer substrates are present, in agreement with the EMSA results.

To further assess the role of protein-protein interactions in cooperative B35SSB-ssDNA binding, we performed cross-linking assays adding ssDNA in excess to the reaction. Although the B35SSB:ssDNA ratio was increased to 1:100, the intensity of the high molecular weight band corresponding to oligomeric species remained unchanged (Figure 4D). This indicates that B35SSB oligomers formation is not only due to the cross-linked interaction between the B35SSB and ssDNA but it is mostly stabilized by protein-protein interactions. This oligomer formation occurs even at very high ssDNA concentration which indicates that the SSB binds in a non-distributive manner where the binding of one single SSB leads to the binding of the next monomer, and so forth, until the ssDNA molecule is completely covered. These and the EMSA results confirm that B35SSB binds to ssDNA in a highly cooperative mechanism where both protein-DNA and protein-protein interactions play a fundamental role.

### DNA binding is mediated by ionic and hydrophobic interactions

As outlined in the Introduction, SSBs mainly bind ssDNA through the nucleotide bases, *via* base-stacking interactions to aromatic residues and cation-stacking contacts [25]. We aimed to identify the main molecular interactions involved in the ssDNA-binding mechanism of B35SSB and, likely, other B35SSB-related proteins. We selected several residues with diverse chemical properties and conservation degrees in order to analyze their contribution to DNA binding. A total of 13 variants were generated (F48A, S49A, Y63A, Y63F, D85A, V87A, Y117A, Y117F, V124A, K130A, Y150A, Y150F, and K156A), but two of them (D85A and Y117A) gave rise to insoluble protein in the inclusion bodies (Figure S2A). This suggests that these modifications have a severe impairment in the structural stability of the protein. Solubility and expression were also impaired, to a lesser extent in the case of Y63A, whose purification resulted in low yield.

The ssDNA-binding ability was strongly impaired for all the obtained variants, except for Y150A and Y150F (Figure 5A). These two variants showed an ssDNA-binding capacity similar to the WT. In turn, F48A, V87A, Y117F, and V124A formed mainly unstable ssDNA-protein complexes detected as a high proportion of quantified smear in EMSA gels (Figure 4A-B). These variants correspond to hydrophobic residues that may be involved in base-stacking/hydrophobic interactions. Also, the ssDNA-binding capacity of Y63A is strongly impaired while the conservative substitution in the conservative Y63F showed a milder defect on DNA binding, which suggests an important role of Y63 in ssDNA binding by base-stacking interactions. Finally, K130A and K156A ssDNA binding is strongly reduced, indicating that ionic interactions can be involved in B35SSB-ssDNA interaction.

**Figure 5.**
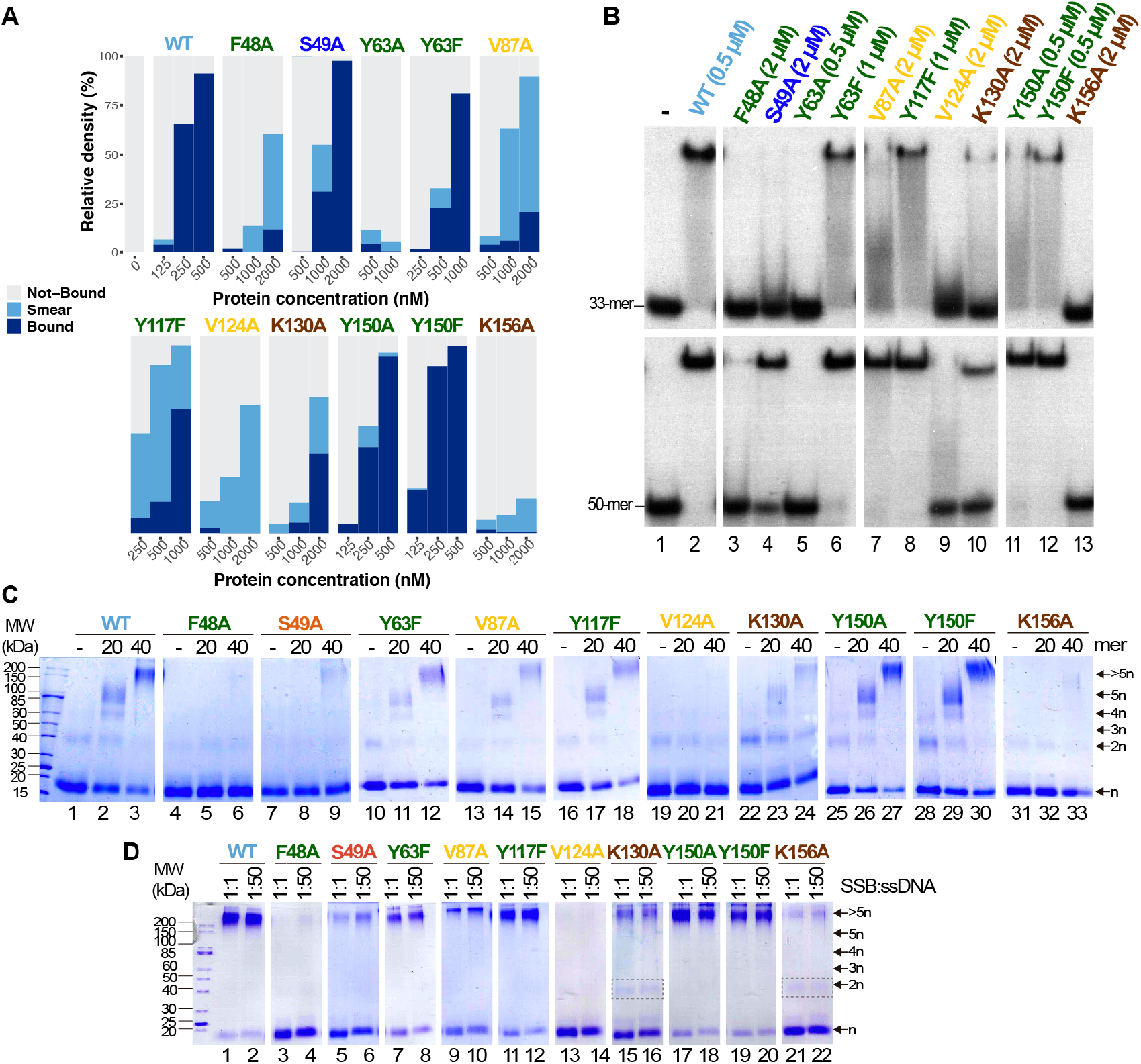
Study of ssDNA-binding ability of B35SSB variants. **(A)** Representative graphic of EMSA assay with B35SSB variants quantified with ImageJ. EMSA was performed using the 50-mer ssDNA probe (Table S1) and the indicated concentration of B35SSB. Note that the B35SSB concentration range was adapted for each variant. **(B)** Effect of ssDNA size in the ssDNA-binding capacity of B35SSB variants. EMSA was performed incubating the indicated concentration of B35SSB or its variants with 33-mer or 50-mer(2) substrates as previously described. **(C)** B35SSB variants cross-linking analysis using different ssDNA sizes. **(D)** Effect of SSB:ssDNA ratio in binding. 1 μM B35SSB (or variants) was incubated either with 1 or 50 μM 50-mer(2) ssDNA, according to the ratio protein:DNA indicated. Dimeric species for K130A and K156A are indicated with a gray rectangle. See Methods for details.

Interestingly, a clear increase in the intensity of protein-DNA complexes was detected for large ssDNA fragments (50-mer) over shorter oligonucleotides (33-mer). As shown in Figure 5B, even the proficient Y150A and Y150F variants formed more stable complexes with the 50-mer substrate, which suggests that Y150 residue may have some role in the interaction with the ssDNA. This effect was stronger for S49A and V87A, although a higher SSB concentration was required to detect stable complexes (lanes 4 and 7). When we use a 33-mer substrate, cooperative binding would be less efficient, hence these results would imply that these variants are not defective in cooperative binding. On the other hand, V124A and F48A could not form stable protein-DNA complexes even with the longer substrate and at high protein concentration (lanes 3 and 9), suggesting a very unstable binding to ssDNA which would not be stabilized in conditions that facilitate cooperative binding. In line with this observation, the use of cross-linking agent could stabilize DNA complexes for Y63F, V87A, Y117F, K130A, Y150A, and Y150F variants (Figure 5C), although only Y150A and Y150F complexes showed a similar intensity to the WT ones. On the other hand, for F48A, S49A, and K156A, oligomers formation was observed as a faint band only when a 40-mer oligo was used as substrate and no oligomers were detected for V124A. These results would mean a severe defect in ssDNA binding, in agreement with the poor binding detected by EMSA.

Finally, the increase of the ssDNA:protein ratio does not affect the results of the cross-linking experiment for any of the analyzed variants (Figure 5D), downplaying a highly specific role in cooperativity. Nevertheless, the loss of charge variants K130A and K156A seems to increase the dimer band, which cannot be detected in the case of the WT and other variants when a large substrate was used, which also may indicate a role of these basic residues in protein-protein interactions and, therefore, in cooperativity.

### A novel, highly divergent OB-fold-related domain in a new group of protein-primed viral SSBs

As mentioned above, protein-primed replicating genomes code for divergent single stranded DNA binding proteins, with non-OB-fold conformation or without structural characterization [46]. The latter is the case of proteins P12 and P19 from PRD1, which are atypical single-stranded DNA binding proteins that would play a key role in the protein-primed viral genome replication, protecting DNA intermediaries and stimulating DNA synthesis [10,47]. These proteins are conserved in PRD1-related viruses, but no ortholog has been detected in other *Tectiviridae* members, including Bam35 [48]. Although no OB-fold protein has been identified in Bam35 proteome, we have showed here that the product of ORF2 is a DNA binding protein with a great specificity for ssDNA, whose binding mechanisms entails base-stacking and ionic interactions. Thus, we can conclude that it shares most of the biochemical properties of OB-fold containing SSBs.

As mentioned above, previous reports had already annotated Bam35 ORF2 product as a putative SSB [34]. In line with this, B35SSB and betatectivirus orthologs are included in a small Pfam family of viral SSBs, along with Φ29 SSB and other proteins (PF17427). We aimed to clarify both the sequence similarity of B35SSB with known SSBs and the presence of a conserved OB-fold. For the first task, we performed Jackhammer [49] iterative searches seeded with the amino acid sequence of B35SSB against the Uniref90 database. We obtained a similar but extended dataset, compared with the PF17427 group. Thus, although we did not detect any sequence corresponding to known OB-fold structures, we obtained several hits corresponding to characterized or predicted SSBs from Φ29 and diverse podoviruses, like GA-1 or asccphi28, other possible phage proteins, as well as diverse protein sequences from bacteria, including *Bacillus sp*. and other *Firmicutes*, which may correspond to misannotated bacteriophage or prophage proteins rather than chromosomal bacterial ORFs. Similar to the Pfam cluster, we also obtained a few metagenomic sequences as well as some eukaryotic proteins from nematodes and insects that might have an origin on horizontal transfer events, though sample contamination cannot be ruled out.

As expected from such a diverse dataset of short sequences, phylogeny reconstruction not always yielded confident clades (Figure 6). However, it should be highlighted that B35SSB and its betatectivirus orthologs form a very compact group, within a well-supported clade (bootstrap value of 98) of more diverse sequences annotated as *Bacillus* proteins, which suggests a single event of acquisition of this sequence in an ancestral betatectivirus.

**Figure 6.**
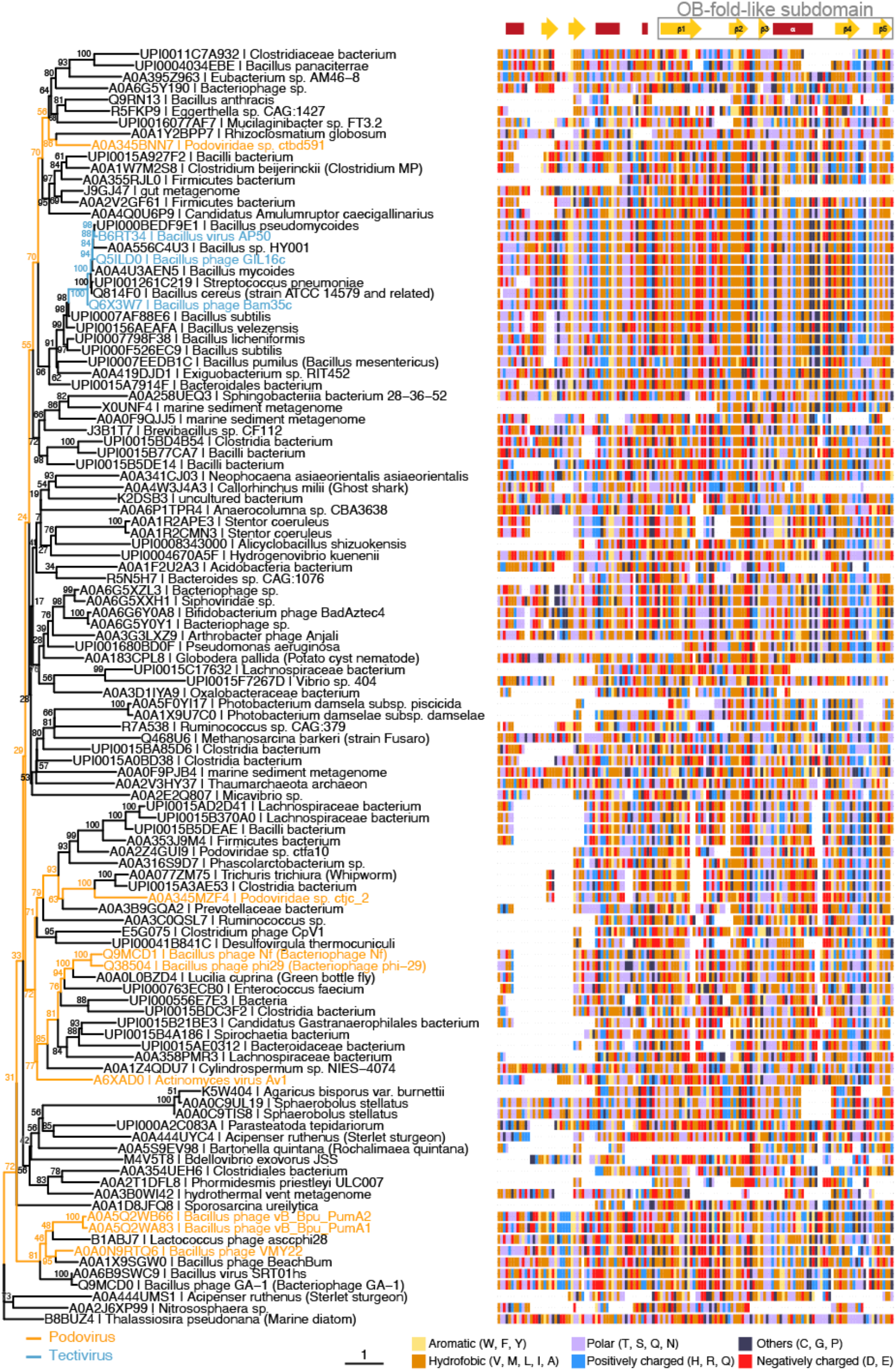
B35SSB and Φ29 related viruses belong to the same clade of SSBs from protein-primed genomes. Maximum-likelihood tree (left) generated using the trimmed multiple sequence alignment of B35SSB-related proteins (right) and visualized with the ggtree package for R software [50]. Strain names of phages belonging to *Picovirinae* (blue) or *Betatectivirus* (orange) groups are indicated. Secondary structure prediction was obtained with the Jpred4 [51] and is indicated above the alignment, with the conserved C-terminal region (residues 76-167 in B35SSB) boxed. Residues are colored by their biochemical properties: aromatic (yellow), hydrophobic (orange), uncharged (purple), positively charged (blue), negatively charged (red), others (dark blue).

The combination of the multiple sequence alignment with secondary structure predictions allowed us to identify a core of about 100 residues that overall share a common fold. Remarkably, the C-terminal part of this fold might be a reminiscent of an OB-fold, as it is made up of five β-strands and one α-helix. Next, we did HHpred search using B35SSB as a query. Among hits, we found the profile of an OB-fold protein (ECOD_001121967_e4m8oA1), albeit with a very low probability (32%) (Figure S4).

To further test the presence of an OB-fold in the Bam35-Φ29 SSBs group, we generated an extended dataset by the addition of sequences from previous phylogenetic analysis of diverse groups of OB-fold-containing cellular and viral SSBs [21,46], updated with our own searches (see Methods), which attempts to span all known SSBs groups. To improve the sensitivity of comparisons, we built HHsearch profiles for all sequences of SSBs, did their all-against-all comparison and clustered the results using CLANS [47] at P-value of 1e-08. Clustering analysis revealed that the new SSBs clade form a compact and independent group, without significant relationship to any particular OB-fold family (Figure S5A). Only when we raised the P-value to 1e-04, connections between the B35-Φ29 SSBs group and other OB-fold groups appeared (Figure S5B), indicating a very remote similarity.

Since diverse sensitive sequence comparisons did not find a clear similarity between the B35-Φ29 SSBs and known OB-folds, we decided to perform structural comparisons. To do so, we first obtained structural models of B35SSB with diverse comparative modeling and contact-based approaches. Our notion that B35SSB has an OB-fold-like domain was supported by three different methods (RaptorX [52], trRossetta [53] and RosettaCM [54,55]), which yielded similar models, of intermediate-good quality, as assessed by VoroMQA and ModFold8 [56,57] (Figure 7A). B35SSB structural models consisted of an N-terminal motif with a disordered region between two α-helixes and a C-terminal domain made up of a five-stranded β-sheet and one α-helix. In agreement with the secondary structure predictions, the structural model of B35SSB somewhat resembles an OB-fold, particularly in the C-terminal two-thirds portion (residues 76-167).

**Figure 7.**
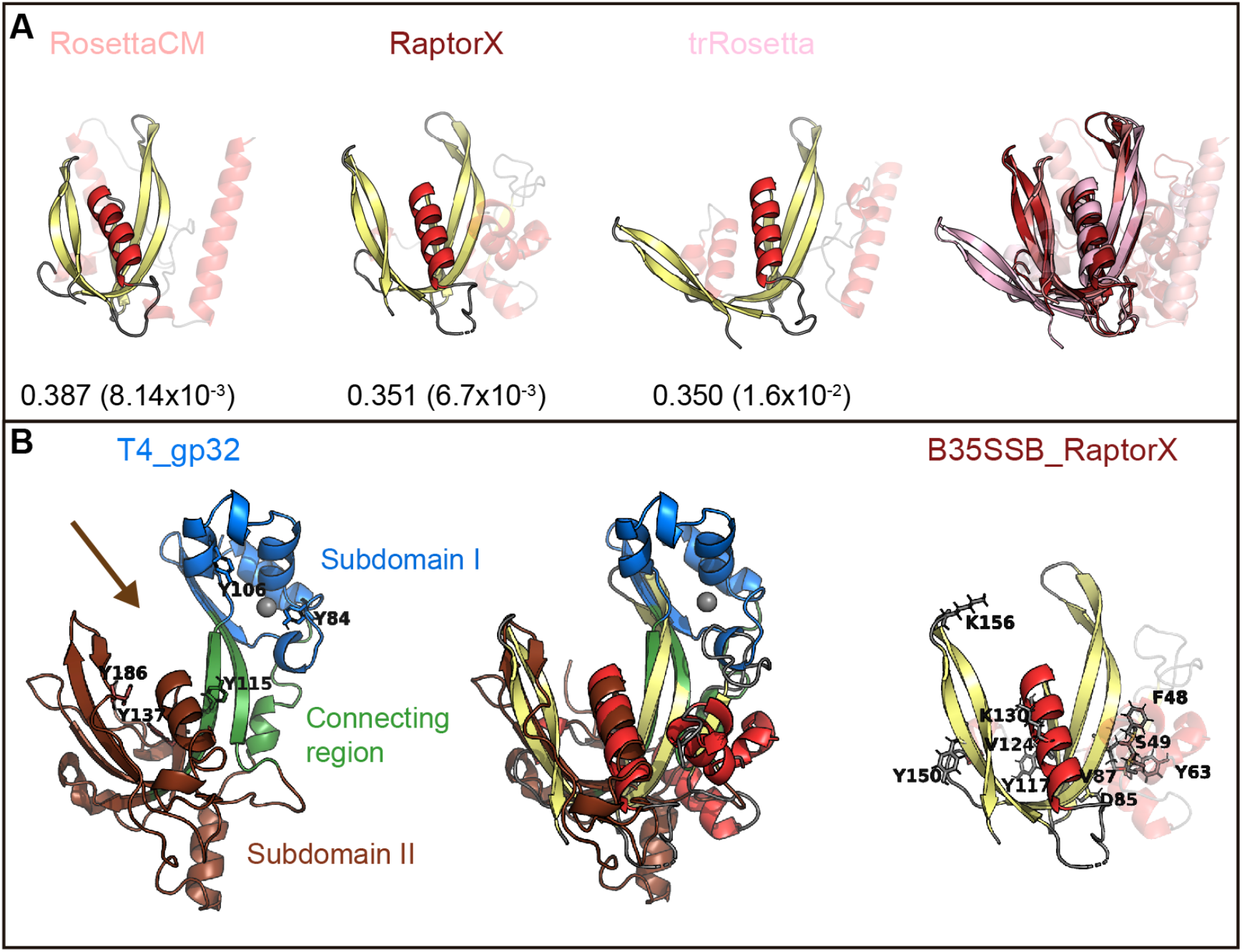
B35SSB and related proteins can constitute a novel SSB fold, divergent to known OB-folds. **(A)** Best structural models of B35SSB, obtained with three different methods as indicated. The N-terminal 76 residues are displayed as semitransparent to show clearly the C-terminal domain rich in β-sheets that might be reminiscent of an OB-fold. The VoroMQA Score and the model confidence E-value from ModFold8 server are also shown. Note that VoroMQA considers good models those with Score >0.4. The rightmost figure represent a structural alignment of the three models. **(B)** Structural alignment of T4 SSB and RaptorX model of B35SSB (middle). Cartoon representation of T4 gp32 (PDB ID 1GPC) is shown on the left. Protein subdomains are colored as in [59] and tyrosine residues involved in DNA binding [60] are highlighted as sticks. The zinc cation bound to T4 gp32 Subdomain I is represented as a grey sphere. The arrow marks the DNA binding cleft. The right image represents the B35SSB model with the residues analyzed displayed as sticks.

We then performed structural searches of the three models using Dali Server [58]. The best hits obtained (Figure S6) included several DNA binding proteins, but only one SSB, the T4 SSB (gp32 protein). The structural alignment between the RaptorX model of B35SSB and the T4 SSB structure [59] (Figure 7B) indicates an overall similarity between the conserved fold of B35-Φ29 SSBs group and the T4 SSB, spanning the β-sheets backbone from subdomains I (Zn finger) and II (DNA binding) and the α-helix that caps the OB-fold.

In conclusion, the conserved betatectiviral SSB contains a predicted alpha-beta complex conserved fold, only distantly related to an OB-fold. This domain seems to be conserved among SSBs from protein-primed replicating viral genomes, spanning Φ29 and related *Picovirinae* proteins, thus constituting the first OB-fold-like protein in a protein-primed genome.

## Discussion

### Extremely cooperative ssDNA binding capacity in B35SSB

The biochemical characterization of B35SSB confirmed some features common to all SSBs, and some specific ones shared only with other protein-primed replication systems. Thus, we found that B35SSB has a high specificity for ssDNA over dsDNA and RNA. Both B35SSB and Φ29SSB display negligible dsDNA-binding capacity [61]. This contrasts with other DNA-binding proteins of protein-primed replicons such as the PRD1 P12 and the adenovirus DBP in which dsDNA-binding ability is reduced but clearly better than in B35SSB [10,62]. Interestingly, bacteriophage T4 SSB also shows a high specificity for ssDNA over dsDNA, due to the narrow DNA binding groove [59], which would have similar width in B35SSB according to RosetaCM and RaptorX models and thus could also explain the negligible binding to dsDNA of B35SSB (Figure 7).

B35SSB binds to ssDNA in a non-sequence dependent manner, which also correlates with the non-specific DNA binding by most SSBs as well as the related pBClin15 ORF 2 gp (85% amino acid identity with B35SSB) [21,33]. B35SSB does show a preference for pyrimidines, which has also been described for some OB-fold proteins and explained by a less thermodynamically favorable interaction of aromatic side chains with the purines due to steric hindrance and inefficient base-stacking [63,64].

Thus, our results point to a DNA-binding mechanism involving base-stacking interactions with aromatic residues, similar to that of OB fold-containing SSBs.

In line with the results of the Y2H Bam35 intraviral interactome [65], where no self-interaction of this protein was observed, B35SSB is a monomer in solution. The podoviral Φ29SSB and Nf SSBs are also monomers but GA-1 SSB is generally a hexamer in solution, although at high salt concentration (200 mM NaCl) it is detected as a monomer [66]. Although a similar salt concentration was used in our ultracentrifugation assays, a lower concentration was used in the crosslinking assays, confirming the monomeric state of B35SSB regardless of the salt concentration.

The B35SSB very high cooperativity was further supported by an enhanced binding to longer ssDNA substrates and, more importantly, by cross-linking assays in which, even at an extremely low B35SSB:ssDNA ratio (1:100), only full-length covered substrates could be detected (Figure 4D). In contrast, *S. solfataricus* SSB monomers can be cross-linked at a ratio of 1:1 but not in the presence of a somewhat molar excess of ssDNA (1:4), indicating that in non-processive binding the monomers distribute evenly along and between the nucleotides [64]. Positive cooperativity can be mediated by protein-protein direct contacts between the nearest neighbors such as *E. coli, T. thermophilus*, and T4 SSBs [67–69], which can be dependent on salt concentration, [40,41]. However, in the case of B35SSB cooperative binding is unaffected by salt concentration (Figure 5B). The results on B35SSB cooperativity show a virtually unlimited cooperativity based on nearest-neighbor interaction, similar to the case of the 35-nt mode of *E. coli* SSB [70]. Comparing B35SSB and EcoSSB, a higher concentration of B35SSB is needed to shift all the 50 nt ssDNA probe, suggesting a smaller binding site. Similar to other monomeric SSBs, B35SSB occludes a small binding site of 4-5 nt [27,71]. This binding site suggests that rather than wrapping around the SSB, the ssDNA is bound initially through an exposed binding surface and B35SSB then multimerizes sequentially along the substrate in a contiguous fashion, as proposed for T4, *S. solfataricus* or Enc34 SSBs [28,43,72]. B35SSB binds ssDNA in a highly cooperative manner, suggesting that both protein-protein and protein-DNA interactions contribute to the formation of stable ssDNA-SSB complexes.

In line with their cooperative binding, a lower concentration of B35SSB than of Φ29SSB is required to obtain proficient DNA binding [27]. This lower affinity in Φ29SSB has been linked to the necessity of the SSB dissociation to allow the DNA polymerase advance during processive replication [73], which would require a fine-tuned SSB-DNA ratio (Figure 4). The SSB of GA-1 is considerably different in sequence, oligomerization, and ssDNA affinity [27,73]. Interestingly, Φ29SSB shares less similarity and identity with GA-1 SSB than GA-1 SSB with B35SSB (17% and 21% identity and 42% and 59% similarity respectively). GA-1 SSB and B35SSB are similar in the N-terminal region but different to Φ29SSB (Figure S7). On the other hand, although the divergence between B35SSB and Φ29SSB and is bigger, the predicted secondary structure of the C-terminal region is very similar. Altogether, we can differentiate two main regions in B35SSB, the C-terminal domain, a highly conserved novel fold that probably plays an essential role in the ssDNA-binding of Bam35-Φ29 SSBs proteins and an N-terminal domain, less conserved and that might also contribute to DNA binding by enhancing the cooperativity, which is compatible with the DNA binding properties of B35SSB variants F48A and S49A.

In some cases, such as GA-1 and T4 SSBs, the N-terminal domain has a role in cooperativity through self-association mediated by electrostatic interactions [66,69]. Nevertheless, within the C-terminal region of B35SSB, we have identified two conserved positively charged residues, K130 and K156, whose non-conservative mutation severely affects the ssDNA-binding ability. EMSA and cross-linking assays suggest a putative role in cooperative binding. However, their predicted location at the limits of the binding groove in the structural models (Figure 7D) would also be compatible with a role in the interactions with the sugar-phosphate backbone. That notwithstanding, it should be emphasized that dissection of a unique role in protein-DNA or protein-protein interactions is challenging, as cooperative binding requires both features. Indeed, high salt concentrations disrupt ssDNA binding with no detected effect on cooperativity, suggesting that electrostatic interactions can be required for both protein-protein or protein-DNA contacts. Another residue located in this region, V124, may also contribute to B35SSB cooperativity. This residue is well conserved among Bam35-related SSBs, its ssDNA-binding ability is highly unstable and could not be stabilized in cross-linking assays where no oligomers were observed. SSBs-ssDNA binding is mainly driven by base-stacking binding to aromatic residues, cation-stacking interactions, and, additionally, hydrophobic and hydrogen-bonding interactions, salt bridges and H bonds (reviewed in ref. 25). The generation of B35SSB variants according to this criterion and the conservation level among either the Bam35-Φ29 SSBs or the group of betatectiviral SSBs helped us to propose some residues essential for cooperativity (discussed above) and ssDNA binding. As expected, the level of conservation is in agreement with the role of the different residues as it is shown in the case of the barely affected variants of the poorly conserved residue Y150 (Figure 5 and Figure S3).

The ssDNA binding groove of T4 SSB contains three aromatic residues (Y115, Y137 and Y186), being Y115 and Y186 involved in ssDNA binding, whereas Y137 seems dispensable [60]. Thus, the protein-DNA interactions within the putative ssDNA binding groove of B35SSB would be different, as Y117 would correspond to Y137. According to our structural models, Y117 would be within the putative ssDNA binding groove (Figure 7B), but contrary to Y137 it is essential for base stacking interactions. Indeed, B35SSB residue Y117 would correspond to the residue Y76 in Φ29SSB (Figure S7). Intrinsic tyrosine fluorescence quenching assays indicated that Φ29SSB Y76 would be involved in the ssDNA binding [38,75]. The phenotype of the conservative replacement of Φ29SSB Y76 to phenylalanine is similar to the obtained with the variant Y117F of B35SSB and the non-conservative replacement to alanine resulted in a loss of solubility in both proteins. Additionally, the Φ29SSB variant Y76S which conserves the OH group but not the aromatic ring resulted in a loss of binding ability [74,76]. Altogether, these results point out this amino acid as an essential residue in ssDNA binding through base-stacking interactions, which is in agreement with its high conservation among the whole clade of viral SSBs.

On the other hand, the B35SSB Y63 variants showed a similar phenotype as Y117 (Figure 5) suggesting a similar role in ssDNA binding. This residue is also well conserved among this cluster as an aromatic residue that can be tyrosine or phenylalanine (Figure S3) which indicates the relevance of the aromatic ring similarly to Y117. However, the instability of the Y63A variant (Figure S2) and its position outside the predicted DNA binding groove (Figure 7), suggests that this residue may be also essential for proper protein folding and/or stability in solution, similar to tyrosines 73 and 92 in the case of T4 SSB [60]. Lastly, among the residues only conserved in betatectiviral SSBs, F48, S49, D85, and V87, our results suggest that F48 could also be necessary for ssDNA binding while D85 and V87 would be required the stability of the protein as their substitution leads to insoluble proteins (Figure S2).

Overall, in the absence of a more detailed structural characterization of the protein, we can conclude that we have identified several aromatic and positively charged conserved residues essential for ssDNA-binding capacity (F48, Y63, Y117, K130, and K156), being aromatic residues essential for protein-DNA interaction.

Besides its specific binding ability to ssDNA, we have characterized the role of B35SSB enhancing processive DNA replication and its DNA unwinding capacity. The stimulation of M13 ssDNA processive replication, a situation that resembles the replicative intermediates generated during Φ29-like DNA replication of linear genomes, has been described for other SSBs such as Φ29, GA-1, and Nf SSBs. In those cases, the SSB prevents the formation of non-productive binding of the DNA polymerase to ssDNA [73]. On the other hand, the helix-destabilizing capacity of T4 and Φ29-like SSBs has been proposed to reduce the energy required for the unwinding process in DNA replication [38,77], which would contribute to the replication stimulation by B35SSB. The unwinding capacity has also been linked to a role in the initiation step of replication [39]. However, we did not detect an effect of B35SSB in the early steps of TP-DNA replication suggesting that its function in replication would occur later during processive replication. On the other hand, it should be noted that the SSBs role in TP-initiation has been described for adenovirus and PRD1 SSBs and has been suggested to be related to, among others, their capacity to bind dsDNA, absent in B35SSB [10,22].

The role of multiple SSBs in replication can be linked to their interaction with other replication proteins, normally polymerases, as illustrated by *E. coli*, T7 and T4 SSBs [26]. Particularly, the increase of the elongation rate and the dissociation of the T7 SSB is associated with an interaction with its cognate DNA polymerase [78]. In the case of B35SSB, *in vitro* stimulation of its cognate DNA polymerase seems unspecific. Together with the absence of detected interactions between the B35SSB and other viral proteins, these results point to a generalized role of B35SSB in replication that does not need specific interactions with other viral factors [65]. This is also the case for Φ29SSB, whose interaction with its cognate polymerase has not been detected and indeed it might be dispensable for viral replication [74,79].

### A highly-divergent OB-fold-like SSB in protein-primed viral genomes

As explained above, oligonucleotide/Oligosaccharide (OB)-fold domains are present in most of single-stranded DNA binding proteins (SSBs) and other DNA binding proteins implicated in genome maintenance and safeguarding in all domains of life. In dsDNA viruses, the OB-fold based SSBs are very common, and they have been classified into five groups, each one represented by the structures of the SSBs of *E. coli*, phages T7 and T4, herpesvirus I3 protein and the archaeo-eukaryotic Replication Protein A [46]. However, the dsDNA viruses with a protein-primed DNA replication mechanism could be one of the very few exceptions to the universal presence OB-fold in SSBs [20]. In this work, we have characterized the product of ORF 2 of Bam35 (B35SSB) as a proficient SSB that would be involved in the protein-primed replication of the viral genome. Phylogenetic analysis and structural predictions indicate that B35SSB shares a common fold with a diverse group of sequences which include unrelated podoviruses with a protein-primed mechanism, such as *Bacillus* viruses Φ29 or GA-1. This fold, covering the 91 C-terminal residues of B35SSB is made up of five β-strands and two α-helixes and thus it may resemble an OB-fold. However, cluster analysis showed that similarity with known OB-fold containing proteins is very remote. Structural comparisons showed a possible similarity with T4 SSB, although it could not be detected with some of the models, underlining the distant relationship of this conserved fold with known OB-fold containing SSBs, as predicted from the clustering results.

Therefore, we can conclude that Bam35-Φ29 SSBs contain a novel and conserved fold, only distantly related to the canonical OB-fold that may have evolved independently. Within this group, B35SSB and betatectiviral orthologs form a highly compact clade within a very diverse group of sequences including Φ29SSB and related viruses from genus *Salasvirus* and related *Picovirinae*, along with predicted proteins from genomic and metagenomic projects that were annotated as bacterial but might as well be encoded by unforeseen viruses. Although other tectiviruses are described or suggested to replicate their genome by a protein-primed mechanism [80–82], B35SSB would be specific for betatectiviruses and, strikingly, clearly related with highly distant podoviruses. In line with this, viruses from *Betatectivirus* and *Salasvirus* genera infect related hosts from *Bacillus* genus [83], which supports a possible ancient common ancestor for Φ29 and Bam35 SSBs, providing a possible scenario for a gene transfer event.

In conclusion, we have identified and thoroughly characterized the single-stranded protein of *Bacillus* virus Bam35, highly conserved among betatectiviruses. This protein belongs to a novel family of SSBs from protein-primed replicating genomes whose conserved fold may be a diverged OB-fold, only distantly related to the T4 group of OB-fold containing SSBs. Thus, Bam35-Φ29 SSBs would represent the first occurrence of an OB-fold-like domain in a protein-primed virus. Within this group, B35SSB stands out by its high cooperativity and ssDNA-binding capacity and may play an important role in DNA stabilization during protein-primed replication.

## Methods

### Proteins

The Bam35 TP (B35TP), Bam35 DNA polymerase (B35DNAP), Φ29 DNA polymerase (Φ29DNAP), Φ29 terminal protein (Φ29TP) and Φ29 P5 (Φ29SSB) were from the laboratory stocks and purified as described [37,61,95,96]. *Escherichia coli* SSB (EcoSSB) was purchased from Qiagen.

The Bam35 genomic DNA was used to amplify the gene 2 flanked by KpnI and BamHI sites by PCR with B35SSB_FW_KpnI and B35sSB_RV_BamHI primers (Table S1) and Vent DNA Polymerase (New England Biolabs). The digested PCR product was cloned into a pET-52b(+) plasmid (Novagen) in frame with an N-terminal Strep-tag II obtaining the B35SSB expression vector pET52b::B35SSB. B35SSB mutants were obtained by direct mutagenesis using different oligonucleotides (Table S1) and Pfu DNA Polymerase (Agilent). The expression vectors were used to transform *E. coli* XL-1 and verified by sequencing the complete ORF2 using T7_FW primer. B35SSB and its variants were expressed in *E. coli* BL21(DE3) harboring the pET52b::B35SSB plasmid. Cultures were grown in 200 ml of ZYM-5052 autoinduction medium [97] with 150 mg/l ampicillin for 16 hours at 26 or 30 °C. Cells were harvested and disrupted by grinding with alumina, suspended in 6 ml of buffer A (50 mM Tris-HCl pH 8, 1 mM EDTA, 0.04% (v/v) β-mercaptoethanol, 500 mM NaCl, cOmplete™ protease inhibitor cocktail (Roche), 5% (v/v) glycerol) per gram of bacteria and sonicated to reduce the viscosity of the extract by fragmentation of the genomic DNA. Samples were centrifuged for 5 min at 480 x g at 4 °C to remove cell debris and alumina. To extract the soluble protein fraction, samples were centrifuged for 20 min at 17,200 x g at 4 °C. Soluble fractions were applied to 1 ml of Strep-tactin column (IBA) per 3 grams of bacterial pellet, according to the manufacturer’s instructions. The resin was washed with 10 column bed volumes (CV) of buffer A and 15 CV of Strep-tactin washing buffer (IBA). Proteins were eluted in 5 CV of Strep-tactin elution buffer (IBA) and dialyzed overnight against 50 mM Tris-HCl pH 7.5, 50% (v/v) glycerol, 1 mM EDTA, 7 mM β-mercaptoethanol and 0.1 M NaCl. Final protein concentration and purity were analyzed by SDS/PAGE followed by Coomassie blue staining.

Although different growing temperatures were assayed for the expression of D85A, Y117A, and Y63A (Figure S2A), only small amounts of soluble proteins were obtained precluding purification and analysis of D85A and Y117A and resulting in low concentration and purity of Y63A.

The Strep-tag II from the B35SSB fusion protein was removed by digestion using HRV 3C Protease. The digestion reaction of 15 μg of B35SSB was carried out in a final volume of 100 μl of digestion buffer (50 mM Tris-HCl pH 7.5, 150 mM NaCl, 1 mM EDTA, 0.05% (v/v) Tween 20, 1 mM DTT) with 3 units of HRV 3C Protease (Novagen) for 16 hours at 4 °C (Figure S2-B, lanes 1-3). To remove the protease, the reaction was incubated with 100 μl of Glutathione Sepharose Fast Flow resin (Sigma-Aldrich) for 1.5 hours at 4 °C on a rotating wheel. After centrifugation for 1 minute at 1,000 x *g* at 4 °C, the supernatant containing the purified untagged protein was collected (lane 6). To evaluate the yield of the purification procedure, the resin was further washed with 50 μl of digestion buffer (lane 4), and the remaining bound protein was eluted by incubation with 20 μl of 2X Laemmli SDS-PAGE sample buffer (8% (v/v) 2-mercaptoethanol, 3.6% (w/v) SDS, 24% (v/v) glycerol, 240 mM Tris HCl pH 6.8, 0.010% (w/v) bromophenol blue) for 3 min at 95 °C (lane 5). We then verified that the N-terminal Strep-tag fusion does not affect either B35SSB oligomeric state (Figure S2-C) or its capacity to stimulate processive DNA replication (Figure S2-D). These results indicate that the presence of the N-terminal Strep-tag does not have an important impact on B35SSB characteristics.

### Analysis of nucleic acid binding capacity

Nucleic acid binding capacity was analyzed by gel mobility shift assays (EMSA). The ssDNA and RNA substrates (Table S1) were 5’ labeled with [γ-^32^P]ATP and T4 polynucleotide kinase. The dsDNA substrate was obtained by hybridization of the labeled 50-mer oligonucleotide with an excess of 50-mer complementary (50-mer_c) unlabeled oligonucleotide in hybridization buffer (0.2 M NaCl, 60 mM Tris-HCl pH 7.5). Binding between the indicated substrate and protein was carried out in a final volume of 20 μl in binding buffer (50 mM Tris-HCl pH 7.5, 1 mM DTT, 4% (v/v) glycerol, 0.1 mg/ml BSA, 0.05% (v/v) Tween 20, 1 mM EDTA, 50 mM NaCl) for 10 min at 4 °C. Samples were analyzed by electrophoresis in 4% (w/v) polyacrylamide gels (acrylamide/bis-acrylamide 80=1, w/w) containing 12 mM Tris-acetate, pH 7.5 and 1 mM EDTA, and run at 4 °C in the same buffer. Protein-DNA complexes were detected by autoradiography as a shift in the migrating position of the labeled DNA. Quantification of protein-DNA complexes was performed using ImageJ software and plots were obtained using the ggplot package for R software [98]

### Multimerization and cooperativity assays

Analysis of oligomerization estate by analytical ultracentrifugaction was carried out by glycerol gradients (15-30%) with 50 mM Tris-HCl, 1 mM EDTA, 7 mM 2-mercaptoethanol and 200 mM NaCl were formed in 5-ml Beckman polyallomer centrifuge tubes (13 x 51 mm). On top of the glycerol gradient, 200 μl of a solution containing 8 μg of B35SSB, and 5 μg of BSA (Sigma-Aldrich) and Φ29TP as molecular weight markers, were loaded. Samples were centrifuged at 58,000 rpm in a Beckman TST 60.4 rotor for 24 hours at 4 °C. Gradients were fractionated from the bottom of the tube and analyzed by SDS– PAGE followed by protein staining with Coomassie Brilliant Blue.

Glutaraldehyde cross-linking assays of B35SSB in presence of different ssDNA substrates (Table S1) were carried out as described [42]. Briefly, B35SSB was incubated with the indicated ssDNA oligonucleotides for 10 minutes at room temperature (RT) in 100 μl final volume of cross-linking buffer (50 mM sodium phosphate buffer pH 7.5, 50 mM NaCl, 0.05% (v/v) Tween 20). Although different substrates and protein:DNA ratio were tested, the standard assay contained 2 μg of B35SSB (1μM) incubated with 165 ng of the indicated oligonucleotide (0.1 μM for a 50-mer substrate).

Cross-links were formed by the addition of 1% (v/v) glutaraldehyde and incubation for 4 min at 20 °C. The reaction was quenched by adding 5 μl of 2 M NaBH4 (freshly prepared in 0.1 M NaOH) for 20 min at 20 °C. Total protein was precipitated by adding 0.03% (v/v) sodium deoxycholate and 3% (v/v) TCA. Samples were incubated in ice for 15 min and centrifugated for 10 min at 14,000 x g at 4 °C. The supernatant was discarded, and the pellet was resuspended in 20 μl of 1X Laemmli SDS-PAGE sample buffer (4% (v/v) 2-mercaptoethanol, 1.8% (w/v) SDS, 12% (v/v) glycerol, 120 mM Tris HCl pH 6.8, 0.005% (w/v) bromophenol blue) and heated to 95 °C for 5 min. Samples were analyzed by electrophoresis in 8-16% polyacrylamide (w/v) gradient SDS gels followed by staining with Coomassie Brilliant Blue.

### Helix-destabilizing assay

Unwinding assays substrate was obtained by hybridization of the genomic M13mp18 single-stranded circular DNA with the labeled primed M13 universal primer (M13UP: 5’ CAGGAAACAGCTATGAC 3’) in hybridization buffer (0.2 M NaCl, 60 mM Tris-HCl). M13UP was labeled with [γ-^32^P]ATP and T4 polynucleotide kinase [99]. The assay was carried out in 12.5 μl of unwinding buffer (50 mM Tris-HCl pH 7.5, 1 mM DTT, 4% (v/v) glycerol, 0.1 mg/ml BSA, 0.05% (v/v) Tween 20, 1 mM EDTA, 50 mM NaCl) with 4 nM labeled primed M13 ssDNA and the indicated concentration of B35SSB or Φ29SSB. Reactions were incubated for 1 hour at 37 °C and stopped with 1.25 μl of 0.25% (w/v) bromophenol blue, 0.25% (w/v) xylene cyanol, 30% (v/v) glycerol and 0.5% (w/v) SDS. Samples were analyzed by electrophoresis in 0.1% (w/v) SDS 8-16% (w/v) gradient polyacrylamide gels. Gels were dried and the helix-destabilizing activity was detected by autoradiography as a change in the mobility of the labeled primer due to its displacement from the M13 DNA molecule.

### Protein-primed initiation and elongation assays

Reactions of TP-initiation and elongation were carried out, as described in [5], in a final volume of 25 μl of reaction buffer (50 mM Tris-HCl pH 8.5, 1 mM DTT, 4% (v/v) glycerol, 0.1 mg/ml BSA, 0.05% (v/v) Tween 20) with 5 μM of dTTP (initiation assays) or all 4 dNTPs (elongation assays) supplemented with 2 μCi of [α-^32^P]dTTP, a preincubated mix (10 min) with 2.5 nM of B35DNAP and 34 nM of B35TP, 10 mM MgCl_2_, 315 nM of the 29 or 60-mer single-stranded oligonucleotide with Bam35 genome left end sequence (Table S1) as the template and the corresponding concentration of SSB.

### Processive replication of singly primed M13 DNA

The primed M13 ssDNA was obtained by hybridization of the genomic M13mp18 single-stranded circular DNA with the M13UP primer in hybridization buffer (0.2 M NaCl, 60 mM Tris-HCl). Reactions of replication were carried out in a final volume of 25 μl of reaction buffer (50 mM Tris-HCl, 1 mM DTT, 4% (v/v) glycerol, 0.1 mg/ml BSA, 0.5% Tween 20) with 50 nM B35DNAP, 1.4 nM primed M13 ssDNA, 40 μM dNTPs, 10 mM MgCl2 and 0.5 μCi [α-^32^P]dATP, and the indicated concentrations of SSB. To increase the efficiency of replication, the polymerase and the template were incubated for 10 minutes at RT prior to the addition of the SSB. Reactions were incubated at 37 °C for the indicated times and stopped by adding 30 mM EDTA and 0.5% (v/v) SDS. Non-incorporated radiolabeled dATP was removed by Sephadex G-50 spin filtration. The lambda DNA ladder was labeled by filling in with Klenow fragment (New England Biolabs) in the presence of [α-^32^P]dATP. Results of replication were analyzed by electrophoresis in alkaline 0.7% agarose gels, and the gels were subsequently dried and autoradiographed.

### B35SSB sequence analysis and phylogeny

Bam35 ORF2 product sequence (B35SSB, UniProt database accession Q6X3W7) was used as the protein query to find related proteins using the Jackhmmer tool from the HMMER web server [49]. The analysis was performed using Uniref90 as database and 10^-5^ as E-value. Search converged after 5 iterations and 112 significant query matches were obtained. The retrieved protein sequences were used to obtain a multiple sequence alignment (MSA) using MAFFT (parameters: -- maxiterate 1000 --retree 1) [44]. The MSA was then trimmed with trimAl (parameter-gappyout) [84] and used to generate a best-fit maximum-likelihood phylogeny using IQTREE2 version 2.0.7 (parameters -B 5000 -alrt 5000 -redo) [85,86]. According to the Bayesian information criterion, the selected best-fit model was LG+F+R5. Lastly, the tree was visualized with the ggtree package for R software [87]. Secondary structure prediction of the MSA was obtained with the JPred4 server through the JalView web server version 2.11 [45,51]. To assess the possible relationship of the Bam35-Φ29 SSBs group with previously known OB-fold containing SSBs, we used an alignment-independent clustering approach. First, a comprehensive dataset aiming to span all known diversity was generated by merging previously used sets of sequences [20,88], updated with our own searches with TopSearch [89] (accessed at https://topsearch.services.came.sbg.ac.at). The final dataset contained 2,750 non-redundant sequences. Then, HMM profiles for each sequence were generated and the profiles were compared all-against-all with HHsearch to cluster them with CLANS [90,91].

The B35SSB structural models were obtained with five different servers (Jan 2021), based either in fold-recognition or distant-based approaches, i-Tasser [92], Phyre2 [93], RaptorX [52], trRossetta [53] and RosettaCM [54,55]. Besides the evaluation provided by each method, all models were assessed by VoroMQA and ModFold8 protein structure quality assessment independent methods [56,57]. Models were then visualized with Open-Source Pymol [94].

Structural searches in PDB database were performed using Dali Protein Structural Comparison Server [58].

## Author Contributions

Conceptualization (MS, MRR), Formal analysis (AL, DK, MRR), Funding Acquisition (MS, MRR), Investigation (AL, DK, MRR), Supervision (MS, MRR), Writing – Original Draft Preparation (AL, MRR), Writing – Review & Edit (AL, DK, MRR).

## Acknowledgments

This work was funded by grants from Spanish Ministry of Science, Innovation and Universities [PGC2018-093726-B-100 AEI/FEDER UE to M.S. and PGC2018-093723-A-100 to M.R.R.] and Fundación Ramón Areces [VirHostOmics]. A. L. was holder of a PhD fellowship [FPU15/05797] from the Spanish Ministry of Science, Innovation and Universities.

An institutional grant from Fundación Ramón Areces and Banco Santander to the Centro de Biología Molecular Severo Ochoa is also acknowledged.

We are also grateful to all the members of the MRR lab for discussions and suggestions.

## References

[1] Gillis A, Fayad N, Makart L, Bolotin A, Sorokin A, Kallassy M, et al. Role of plasmid plasticity and mobile genetic elements in the entomopathogen Bacillus thuringiensis serovar israelensis. FEMS Microbiology Reviews 2018;42:829–56. https://doi.org/10.1093/femsre/fuy034.

[2] Ackermann HW, Roy R, Martin M, Murthy MR, Smirnoff WA. Partial characterization of a cubic Bacillus phage. Canadian Journal of Microbiology 1978;24:986–93.

[3] Krupovic M, Koonin EV. Polintons: a hotbed of eukaryotic virus, transposon and plasmid evolution. Nat Rev Microbiol 2015;13:105–15. https://doi.org/10.1038/nrmicro3389.

[4] Koonin EV, Krupovic M, Yutin N. Evolution of double-stranded DNA viruses of eukaryotes: from bacteriophages to transposons to giant viruses. Ann N Y Acad Sci 2015;1341:10–24. https://doi.org/10.1111/nyas.12728.

[5] Berjón-Otero M, Villar L, Salas M, Redrejo-Rodríguez M. Disclosing early steps of protein-primed genome replication of the Gram-positive tectivirus Bam35. Nucleic Acids Res 2016;44:9733–44. https://doi.org/10.1093/nar/gkw673.

[6] Salas M, de Vega M. Protein-Primed Replication of Bacteriophage Φ29 DNA. In: Laurie SK, Marcos Túlio O, editors. The Enzymes, vol. Volume 39, Academic Press; 2016, p. 137–67.

[7] Salas M, Holguera I, Redrejo-Rodriguez M, de Vega M. DNA-binding proteins essential for protein-primed bacteriophage Φ29 DNA replication. Front Mol Biosci 2016;3:37. https://doi.org/10.3389/fmolb.2016.00037.

[8] Savilahti H, Caldentey J, Lundstrom K, Syvaoja JE, Bamford DH. Overexpression, purification, and characterization of *Escherichia coli* bacteriophage PRD1 DNA polymerase. In vitro synthesis of full-length PRD1 DNA with purified proteins. J Biol Chem 1991;266:18737–44.

[9] Blanco L, Lazaro JM, de Vega M, Bonnin A, Salas M. Terminal protein-primed DNA amplification. Proc Natl Acad Sci U S A 1994;91:12198–202.

[10] Pakula TM, Caldentey J, Serrano M, Gutiérrez C, Hermoso JM, Salas M, et al. Characterization of a DNA binding protein of bacteriophage PRD1 involved in DNA replication. Nucleic Acids Res 1990;18:6553–7.

[11] Salas M. Protein-priming of DNA replication. Annual Review of Biochemistry 1991;60:39–71. https://doi.org/10.1146/annurev.bi.60.070191.000351.

[12] Pestryakov PE, Lavrik OI. Mechanisms of single-stranded DNA-binding protein functioning in cellular DNA metabolism. Biochemistry Mosc 2008;73:1388–404. https://doi.org/10.1134/s0006297908130026.

[13] Naue N, Beerbaum M, Bogutzki A, Schmieder P, Curth U. The helicase-binding domain of Escherichia coli DnaG primase interacts with the highly conserved C-terminal region of single-stranded DNA-binding protein. Nucleic Acids Res 2013;41:4507–17. https://doi.org/10.1093/nar/gkt107.

[14] Hernandez AJ, Richardson CC. Gp2.5, the multifunctional bacteriophage T7 single-stranded DNA binding protein. Semin Cell Dev Biol 2019;86:92–101. https://doi.org/10.1016/j.semcdb.2018.03.018.

[15] Byrne BM, Oakley GG. Replication protein A, the laxative that keeps DNA regular: The importance of RPA phosphorylation in maintaining genome stability. Semin Cell Dev Biol 2019;86:112–20. https://doi.org/10.1016/j.semcdb.2018.04.005.

[16] Kur J, Olszewski M, Dlugolecka A, Filipkowski P. Single-stranded DNA-binding proteins (SSBs) -- sources and applications in molecular biology. Acta Biochim Pol 2005;52:569–74.

[17] Perales C, Cava F, Meijer WJ, Berenguer J. Enhancement of DNA, cDNA synthesis and fidelity at high temperatures by a dimeric single-stranded DNA-binding protein. Nucleic Acids Res 2003;31:6473–80.

[18] Chisty LT, Quaglia D, Webb MR. Fluorescent single-stranded DNA-binding protein from Plasmodium falciparum as a biosensor for single-stranded DNA. PLoS ONE 2018;13:e0193272. https://doi.org/10.1371/journal.pone.0193272.

[19] Hyland EM, Rezende LF, Richardson CC. The DNA binding domain of the gene 2.5 single-stranded DNA-binding protein of bacteriophage T7. J Biol Chem 2003;278:7247–56. https://doi.org/10.1074/jbc.M210605200.

[20] Kazlauskas D, Venclovas C. Two distinct SSB protein families in nucleo-cytoplasmic large DNA viruses. Bioinformatics 2012;28:3186–90. https://doi.org/10.1093/bioinformatics/bts626.

[21] Murzin AG. OB(oligonucleotide/oligosaccharide binding)-fold: common structural and functional solution for non-homologous sequences. EMBO J 1993;12:861–7.

[22] Tucker PA, Tsernoglou D, Tucker AD, Coenjaerts FE, Leenders H, van der Vliet PC. Crystal structure of the adenovirus DNA binding protein reveals a hook-on model for cooperative DNA binding. EMBO J 1994;13:2994–3002.

[23] Paytubi S, McMahon SA, Graham S, Liu H, Botting CH, Makarova KS, et al. Displacement of the canonical single-stranded DNA-binding protein in the Thermoproteales. Proceedings of the National Academy of Sciences 2012;109:E398–405. https://doi.org/10.1073/pnas.1113277108.

[24] Boon M, De Zitter E, De Smet J, Wagemans J, Voet M, Pennemann FL, et al. “Drc”, a structurally novel ssDNA-binding transcription regulator of N4-related bacterial viruses. Nucleic Acids Res 2020;48:445–59. https://doi.org/10.1093/nar/gkz1048.

[25] Dickey TH, Altschuler SE, Wuttke DS. Single-stranded DNA-binding proteins: multiple domains for multiple functions. Structure 2013;21:1074–84. https://doi.org/10.1016/j.str.2013.05.013.

[26] Pestryakov PE, Lavrik OI. Mechanisms of single-stranded DNA-binding protein functioning in cellular DNA metabolism. Biochemistry Moscow 2008;73:1388–404. https://doi.org/10.1134/S0006297908130026.

[27] Gascón I, Gutiérrez C, Salas M. Structural and functional comparative study of the complexes formed by viral Φ29, Nf and GA-1 SSB proteins with DNA. J Mol Biol 2000;296:989–99. https://doi.org/10.1006/jmbi.2000.3521.

[28] Jose D, Weitzel SE, Baase WA, von Hippel PH. Mapping the interactions of the single-stranded DNA binding protein of bacteriophage T4 (gp32) with DNA lattices at single nucleotide resolution: gp32 monomer binding. Nucleic Acids Res 2015;43:9276–90. https://doi.org/10.1093/nar/gkv817.

[29] Dubiel K, Myers AR, Kozlov AG, Yang O, Zhang J, Ha T, et al. Structural Mechanisms of Cooperative DNA Binding by Bacterial Single-Stranded DNA-Binding Proteins. J Mol Biol 2019;431:178–95. https://doi.org/10.1016/j.jmb.2018.11.019.

[30] Kim YT, Tabor S, Bortner C, Griffith JD, Richardson CC. Purification and characterization of the bacteriophage T7 gene 2.5 protein. A single-stranded DNA-binding protein. J Biol Chem 1992;267:15022–31.

[31] Kumaran S, Kozlov AG, Lohman TM. Saccharomyces cerevisiae replication protein A binds to single-stranded DNA in multiple salt-dependent modes. Biochemistry 2006;45:11958–73. https://doi.org/10.1021/bi060994r.

[32] Antony E, Lohman TM. Dynamics of E. coli single stranded DNA binding (SSB) protein-DNA complexes. Semin Cell Dev Biol 2019;86:102–11. https://doi.org/10.1016/j.semcdb.2018.03.017.

[33] Stabell FB, Egge-Jacobsen W, Risoen PA, Kolsto AB, Okstad OA. ORF 2 from the *Bacillus cereus* linear plasmid pBClin15 encodes a DNA binding protein. Lett Appl Microbiol 2009;48:51–7. https://doi.org/10.1111/j.1472-765X.2008.02483.x.

[34] Jalasvuori M, Palmu S, Gillis A, Kokko H, Mahillon J, Bamford JK, et al. Identification of five novel tectiviruses in *Bacillus* strains: analysis of a highly variable region generating genetic diversity. Res Microbiol 2013;164:118–26. https://doi.org/10.1016/j.resmic.2012.10.011.

[35] Shereda RD, Kozlov AG, Lohman TM, Cox MM, Keck JL. SSB as an organizer/mobilizer of genome maintenance complexes. Crit Rev Biochem Mol Biol 2008;43:289–318. https://doi.org/10.1080/10409230802341296.

[36] Shi H, Zhang Y, Zhang G, Guo J, Zhang X, Song H, et al. Systematic functional comparative analysis of four single-stranded DNA-binding proteins and their affection on viral RNA metabolism. PLoS ONE 2013;8:e55076. https://doi.org/10.1371/journal.pone.0055076.

[37] Berjón-Otero M, Villar L, de Vega M, Salas M, Redrejo-Rodríguez M. DNA polymerase from temperate phage Bam35 is endowed with processive polymerization and abasic sites translesion synthesis capacity. Proc Natl Acad Sci USA 2015;112:E3476–84. https://doi.org/10.1073/pnas.1510280112.

[38] Soengas MS, Gutiérrez C, Salas M. Helix-destabilizing activity of Φ29 single-stranded DNA binding protein: effect on the elongation rate during strand displacement DNA replication. Journal of Molecular Biology 1995;253:517–29. https://doi.org/10.1006/jmbi.1995.0570.

[39] Matsumoto K, Ishimi Y. Single-stranded-DNA-binding protein-dependent DNA unwinding of the yeast ARS1 region. Molecular and Cellular Biology 1994;14:4624–32. https://doi.org/10.1128/MCB.14.7.4624.

[40] Lohman TM, Overman LB, Datta S. Salt-dependent changes in the DNA binding co-operativity of Escherichia coli single strand binding protein. Journal of Molecular Biology 1986;187:603–15. https://doi.org/10.1016/0022-2836(86)90338-4.

[41] Pant K, Anderson B, Perdana H, Malinowski MA, Win AT, Pabst C, et al. The role of the C-domain of bacteriophage T4 gene 32 protein in ssDNA binding and dsDNA helix-destabilization: Kinetic, single-molecule, and cross-linking studies. PLOS ONE 2018;13:e0194357. https://doi.org/10.1371/journal.pone.0194357.

[42] Jaenicke R, Rudolph R. [12]Refolding and association of oligomeric proteins. Methods in Enzymology, vol. 131, Academic Press; 1986, p. 218–50. https://doi.org/10.1016/0076-6879(86)31043-7.

[43] Gamsjaeger R, Kariawasam R, Gimenez AX, Touma C, McIlwain E, Bernardo RE, et al. The structural basis of DNA binding by the single-stranded DNA-binding protein from Sulfolobus solfataricus. Biochemical Journal 2015;465:337–46. https://doi.org/10.1042/BJ20141140.

[44] Katoh K, Standley DM. MAFFT multiple sequence alignment software version 7: improvements in performance and usability. Mol Biol Evol 2013;30:772–80. https://doi.org/10.1093/molbev/mst010.

[45] Procter JB, Carstairs GM, Soares B, Mourão K, Ofoegbu TC, Barton D, et al. Alignment of Biological Sequences with Jalview. Methods Mol Biol 2021;2231:203–24. https://doi.org/10.1007/978-1-0716-1036-7_13.

[46] Kazlauskas D, Krupovic M, Venclovas C. The logic of DNA replication in double-stranded DNA viruses: insights from global analysis of viral genomes. Nucleic Acids Res 2016;44:4551–64. https://doi.org/10.1093/nar/gkw322.

[47] Pakula TM, Caldentey J, Gutiérrez C, Olkkonen VM, Salas M, Bamford DH. Overproduction, purification, and characterization of DNA-binding protein P19 of bacteriophage PRD1. Gene 1993;126:99–104.

[48] Ravantti JJ, Gaidelyte A, Bamford DH, Bamford JK. Comparative analysis of bacterial viruses Bam35, infecting a gram-positive host, and PRD1, infecting gram-negative hosts, demonstrates a viral lineage. Virology 2003;313:401–14. https://doi.org/S0042682203002952 [pii].

[49] Potter SC, Luciani A, Eddy SR, Park Y, Lopez R, Finn RD. HMMER web server: 2018 update. Nucleic Acids Res 2018;46:W200–4. https://doi.org/10.1093/nar/gky448.

[50] Yu G, Lam TT-Y, Zhu H, Guan Y. Two Methods for Mapping and Visualizing Associated Data on Phylogeny Using *Ggtree*. Molecular Biology and Evolution 2018;35:3041–3. https://doi.org/10.1093/molbev/msy194.

[51] Drozdetskiy A, Cole C, Procter J, Barton GJ. JPred4: a protein secondary structure prediction server. Nucleic Acids Res 2015;43:W389–394. https://doi.org/10.1093/nar/gkv332.

[52] Källberg M, Wang H, Wang S, Peng J, Wang Z, Lu H, et al. Template-based protein structure modeling using the RaptorX web server. Nat Protoc 2012;7:1511–22. https://doi.org/10.1038/nprot.2012.085.

[53] Yang J, Anishchenko I, Park H, Peng Z, Ovchinnikov S, Baker D. Improved protein structure prediction using predicted interresidue orientations. PNAS 2020;117:1496–503. https://doi.org/10.1073/pnas.1914677117.

[54] Kim DE, Chivian D, Baker D. Protein structure prediction and analysis using the Robetta server. Nucleic Acids Res 2004;32:W526–31. https://doi.org/10.1093/nar/gkh468.

[55] Song Y, DiMaio F, Wang RY-R, Kim D, Miles C, Brunette T, et al. High-resolution comparative modeling with RosettaCM. Structure 2013;21:1735–42. https://doi.org/10.1016/j.str.2013.08.005.

[56] Olechnovič K, Venclovas Č. VoroMQA: Assessment of protein structure quality using interatomic contact areas. Proteins 2017;85:1131–45. https://doi.org/10.1002/prot.25278.

[57] Maghrabi AHA, McGuffin LJ. ModFOLD6: an accurate web server for the global and local quality estimation of 3D protein models. Nucleic Acids Res 2017;45:W416–21. https://doi.org/10.1093/nar/gkx332.

[58] Holm L. DALI and the persistence of protein shape. Protein Sci 2020;29:128–40. https://doi.org/10.1002/pro.3749.

[59] Shamoo Y, Friedman AM, Parsons MR, Konigsberg WH, Steitz TA. Crystal structure of a replication fork single-stranded DNA binding protein (T4 gp32) complexed to DNA. Nature 1995;376:362–6. https://doi.org/10.1038/376362a0.

[60] Shamoo Y, Ghosaini LR, Keating KM, Williams KR, Sturtevant JM, Konigsberg WH. Site-specific mutagenesis of T4 gene 32: the role of tyrosine residues in protein-nucleic acid interactions. Biochemistry 1989;28:7409–17. https://doi.org/10.1021/bi00444a039.

[61] Martin G, Salas M. Characterization and cloning of gene 5 of Bacillus subtilis phage phi 29. Gene 1988;67:193–201.

[62] Stuiver MH, van der Vliet PC. Adenovirus DNA-binding protein forms a multimeric protein complex with double-stranded DNA and enhances binding of nuclear factor I. J Virol 1990;64:379–86.

[63] Kozlov AG, Lohman TM. Adenine base unstacking dominates the observed enthalpy and heat capacity changes for the Escherichia coli SSB tetramer binding to single-stranded oligoadenylates. Biochemistry 1999;38:7388–97. https://doi.org/10.1021/bi990309z.

[64] Touma C, Kariawasam R, Gimenez AX, Bernardo RE, Ashton NW, Adams MN, et al. A structural analysis of DNA binding by hSSB1 (NABP2/OBFC2B) in solution. Nucleic Acids Res 2016;44:7963–73. https://doi.org/10.1093/nar/gkw617.

[65] Berjón-Otero M, Lechuga A, Mehla J, Uetz P, Salas M, Redrejo-Rodríguez M. Bam35 tectivirus intraviral interaction map unveils new function and localization of phage ORFan proteins. J Virol 2017;91. https://doi.org/10.1128/JVI.00870-17.

[66] Gascón I, Carrascosa JL, Villar L, Lázaro JM, Salas M. Importance of the N-terminal region of the phage GA-1 single-stranded DNA-binding protein for its self-interaction ability and functionality. J Biol Chem 2002;277:22534–40. https://doi.org/10.1074/jbc.M202430200.

[67] Raghunathan S, Ricard CS, Lohman TM, Waksman G. Crystal structure of the homo-tetrameric DNA binding domain of Escherichia coli single-stranded DNA-binding protein determined by multiwavelength x-ray diffraction on the selenomethionyl protein at 2.9-A resolution. Proc Natl Acad Sci U S A 1997;94:6652–7. https://doi.org/10.1073/pnas.94.13.6652.

[68] Witte G, Fedorov R, Curth U. Biophysical analysis of Thermus aquaticus single-stranded DNA binding protein. Biophys J 2008;94:2269–79. https://doi.org/10.1529/biophysj.107.121533.

[69] Casas-Finet JR, Fischer KR, Karpel RL. Structural basis for the nucleic acid binding cooperativity of bacteriophage T4 gene 32 protein: the (Lys/Arg)3(Ser/Thr)2 (LAST) motif. Proc Natl Acad Sci U S A 1992;89:1050–4. https://doi.org/10.1073/pnas.89.3.1050.

[70] Lohman TM, Ferrari ME. Escherichia coli single-stranded DNA-binding protein: multiple DNA-binding modes and cooperativities. Annu Rev Biochem 1994;63:527–70. https://doi.org/10.1146/annurev.bi.63.070194.002523.

[71] Wadsworth RI, White MF. Identification and properties of the crenarchaeal single-stranded DNA binding protein from Sulfolobus solfataricus. Nucleic Acids Res 2001;29:914–20. https://doi.org/10.1093/nar/29.4.914.

[72] Cernooka E, Rumnieks J, Tars K, Kazaks A. Structural Basis for DNA Recognition of a Single-stranded DNA-binding Protein from Enterobacter Phage Enc34. Sci Rep 2017;7:15529. https://doi.org/10.1038/s41598-017-15774-y.

[73] Gascón I, Lázaro JM, Salas M. Differential functional behavior of viral Φ29, Nf and GA-1 SSB proteins. Nucleic Acids Res 2000;28:2034–42.

[74] de la Torre I, Quiñones V, Salas M, Del Prado A. Tyrosines involved in the activity of phi29 single-stranded DNA binding protein. PLoS ONE 2019;14:e0217248. https://doi.org/10.1371/journal.pone.0217248.

[75] Soengas MS, Esteban JA, Salas M, Gutiérrez C. Complex formation between phage Φ29 single-stranded DNA binding protein and DNA. Journal of Molecular Biology 1994;239:213–26.

[76] Soengas MS. Caracterización estructural y funcional de la proteína de unión a DNA de cadena sencilla del bacteriófago Φ29. PhD Thesis. Universidad Autónoma de Madrid, 1996.

[77] Pant K, Karpel RL, Rouzina I, Williams MC. Mechanical Measurement of Single-molecule Binding Rates: Kinetics of DNA Helix-destabilization by T4 Gene 32 Protein. Journal of Molecular Biology 2004;336:851–70. https://doi.org/10.1016/j.jmb.2003.12.025.

[78] Cerrón F, de Lorenzo S, Lemishko KM, Ciesielski GL, Kaguni LS, Cao FJ, et al. Replicative DNA polymerases promote active displacement of SSB proteins during lagging strand synthesis. Nucleic Acids Res 2019;47:5723–34. https://doi.org/10.1093/nar/gkz249.

[79] Tone T, Takeuchi A, Makino O. Functional linkages between replication proteins of genes 1, 3 and 5 of Bacillus subtilis phage Φ29. Genes Genet Syst 2012;87:347–56.

[80] Caldentey J, Blanco L, Savilahti H, Bamford DH, Salas M. *In vitro* replication of bacteriophage PRD1 DNA. Metal activation of protein-primed initiation and DNA elongation. Nucleic Acids Res 1992;20:3971–6.

[81] Caruso SM, deCarvalho TN, Huynh A, Morcos G, Kuo N, Parsa S, et al. A Novel Genus of Actinobacterial Tectiviridae. Viruses 2019;11. https://doi.org/10.3390/v11121134.

[82] Philippe C, Krupovic M, Jaomanjaka F, Claisse O, Petrel M, le Marrec C. Bacteriophage GC1, a Novel Tectivirus Infecting Gluconobacter Cerinus, an Acetic Acid Bacterium Associated with Wine-Making. Viruses 2018;10. https://doi.org/10.3390/v10010039.

[83] Alcaraz LD, Moreno-Hagelsieb G, Eguiarte LE, Souza V, Herrera-Estrella L, Olmedo G. Understanding the evolutionary relationships and major traits of Bacillus through comparative genomics. BMC Genomics 2010;11:332. https://doi.org/10.1186/1471-2164-11-332.

[84] Capella-Gutierrez S, Silla-Martinez JM, Gabaldon T. trimAl: a tool for automated alignment trimming in large-scale phylogenetic analyses. Bioinformatics 2009;25:1972–3. https://doi.org/10.1093/bioinformatics/btp348.

[85] Kalyaanamoorthy S, Minh BQ, Wong TKF, von Haeseler A, Jermiin LS. ModelFinder: fast model selection for accurate phylogenetic estimates. Nat Methods 2017;14:587–9. https://doi.org/10.1038/nmeth.4285.

[86] Minh BQ, Schmidt HA, Chernomor O, Schrempf D, Woodhams MD, von Haeseler A, et al. IQ-TREE 2: New Models and Efficient Methods for Phylogenetic Inference in the Genomic Era. Molecular Biology and Evolution 2020;37:1530–4. https://doi.org/10.1093/molbev/msaa015.

[87] Yu G. Using ggtree to Visualize Data on Tree-Like Structures. Current Protocols in Bioinformatics 2020;69:e96. https://doi.org/10.1002/cpbi.96.

[88] Szczepankowska AK, Prestel E, Mariadassou M, Bardowski JK, Bidnenko E. Phylogenetic and complementation analysis of a single-stranded DNA binding protein family from lactococcal phages indicates a non-bacterial origin. PLoS ONE 2011;6:e26942. https://doi.org/10.1371/journal.pone.0026942.

[89] Wiederstein M, Gruber M, Frank K, Melo F, Sippl MJ. Structure-based characterization of multiprotein complexes. Structure 2014;22:1063–70. https://doi.org/10.1016/j.str.2014.05.005.

[90] Frickey T, Lupas A. CLANS: a Java application for visualizing protein families based on pairwise similarity. Bioinformatics 2004;20:3702–4. https://doi.org/10.1093/bioinformatics/bth444.

[91] Soding J, Biegert A, Lupas AN. The HHpred interactive server for protein homology detection and structure prediction. Nucleic Acids Res 2005;33:W244–8. https://doi.org/10.1093/nar/gki408.

[92] Roy A, Kucukural A, Zhang Y. I-TASSER: a unified platform for automated protein structure and function prediction. Nat Protoc 2010;5:725–38. https://doi.org/nprot.2010.5 [pii] 10.1038/nprot.2010.5.

[93] Kelley LA, Mezulis S, Yates CM, Wass MN, Sternberg MJ. The Phyre2 web portal for protein modeling, prediction and analysis. Nat Protoc 2015;10:845–58. https://doi.org/10.1038/nprot.2015.053.

[94] Schrödinger, LLC. The PyMOL Molecular Graphics System, Version 2.4 2021.

[95] Lázaro JM, Blanco L, Salas M. Purification of bacteriophage Φ29 DNA polymerase. Methods in Enzymology 1995;262:42–9.

[96] Mencía M, Gella P, Camacho A, de Vega M, Salas M. Terminal protein-primed amplification of heterologous DNA with a minimal replication system based on phage Φ29. Proc Natl Acad Sci USA 2011;108:18655–60. https://doi.org/1114397108 [pii] 10.1073/pnas.1114397108.

[97] Studier FW. Protein production by auto-induction in high density shaking cultures. Protein Expression and Purification 2005;41:207–34.

[98] Wickham H. ggplot2: Elegant Graphics for Data Analysis. New York: Springer-Verlag; 2009. https://doi.org/10.1007/978-0-387-98141-3.

[99] Sambrook J, Russell DW. Molecular cloning: a laboratory manual. 4th ed. New York: Cold Spring Harbor Laboratory Press; 2001.

