## Supplementary Figures and Tables for "Unlimited cooperativity of *Betatectivirus* SSB, a novel DNA binding protein related to an atypical group of SSBs from protein-primed replicating bacterial viruses"

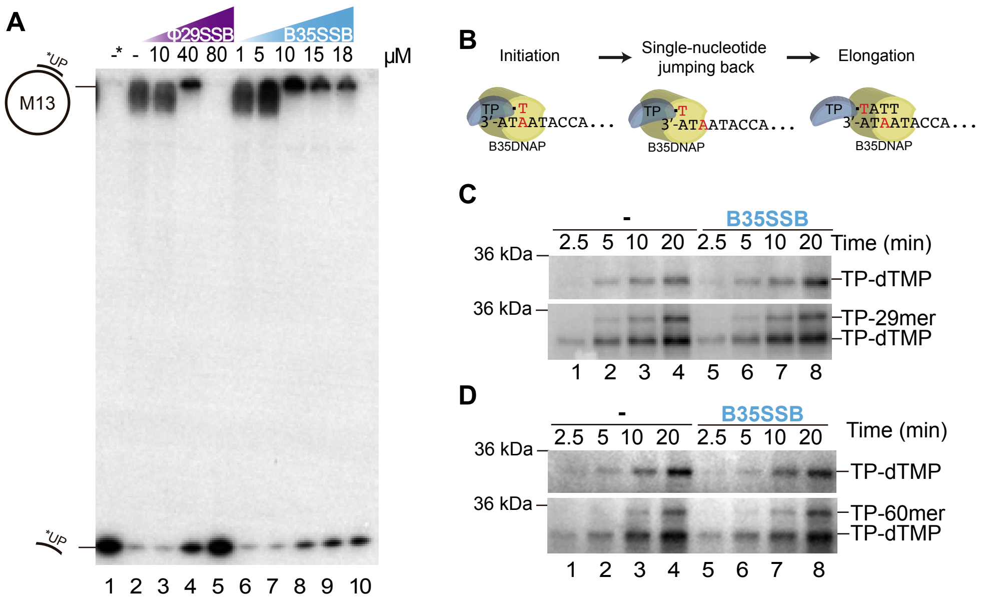


**Figure S1. B35SSB helix-destabilizing capacity and role in early steps of TP-primed DNA replication.**

(A) Helix-destabilizing activity of B35SSB. An M13 ssDNA molecule hybridized to the 5′-labeled M13 UP was incubated with increasing concentrations of either Φ29SSB or B35SSB. After 1 hour at 37 ºC, reactions were stopped and fractionated on a polyacrylamide gel. Positions of the hybrid UP*- M13 and the displaced oligonucleotide are indicated. "-" is the control without protein and "-*" is the control of the heat-denatured substrate.

(B) Schematic representation of Bam35 initial steps of replication. Bam35 protein-primed mechanism can be initiated by the TP deoxythymidylation at conserved tyrosine 194 which is directed by the third base of the template strand. Subsequently, a third-to-first template base single-nucleotide jumping-back process occurs to recover the information of the 3′ end sequence and start the elongation step. Figure adapted from ref. [37].

(C) and (D) SDS-PAGE analysis of Bam35 TP (B35TP)-mediated initiation (with dTTP) and initiation followed by full-length protein-primed replication (with all 4 dNTPs) assays, after the indicated times of reaction. Initiation (upper gel) and replication (bottom gel) reactions with a 29-mer (C) or 60-mer (D) single-stranded oligonucleotide template containing the Bam35 genome origin sequence (Table S1) were performed using B35DNAP, B35TP as primer and 1 μM of B35SSB or no SSB (dash). TP-dTMP initiation complexes and TP-DNA elongation products (TP-29mer or TP-60mer) are indicated. See Methods for details.

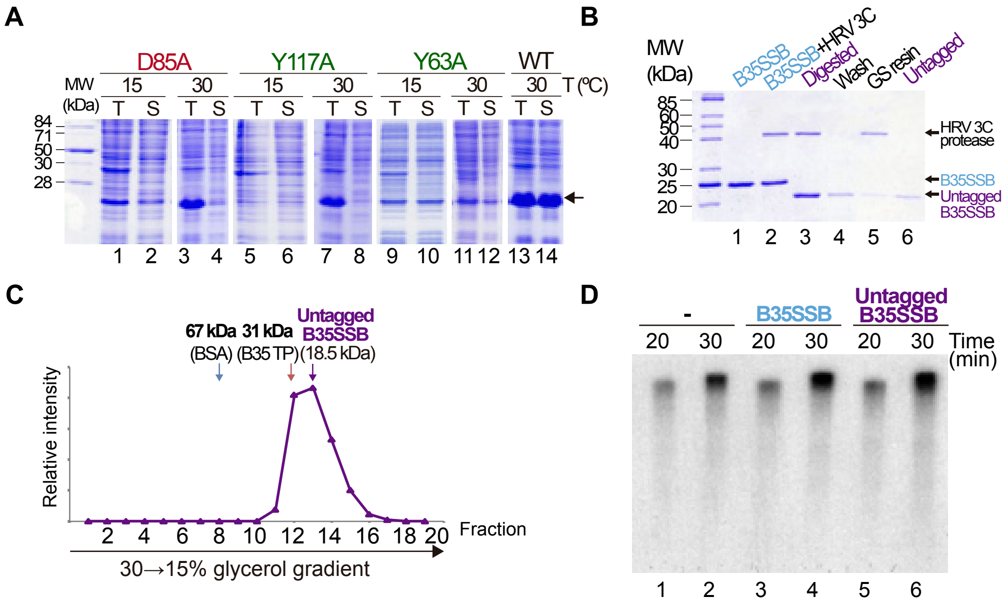


**Figure S2. Overexpression and solubility of B35SSB variants and Strep-tag removal.**

(A) Analysis of expression and solubility levels of D85A, Y117A and Y63A variants and WT at different temperatures. Cultures were grown overnight at 15 or 30ºC and samples corresponding to the total protein extract (T) and the soluble fraction (S) obtained after high-speed centrifugation for 10 minutes were subjected to electrophoresis in 15% (w/v) SDS polyacrylamide gels and visualized by Coomassie staining.
(B) SDS-PAGE visualized by Coomassie staining of the B35SSB Strep-tag cleavage and subsequent purification of the untagged protein. The untagged B35SSB was obtained by Strep-tag cleavage using HRV 3C Protease. The resulting digested protein was purified using Glutathione Sepharose Fast Flow (GS) resin (Sigma-Aldrich).
(C) Determination of the oligomerization state of the untagged B35SSB in solution. Untagged B35SSB (8 µg) was subjected to sedimentation in 15-30% (w/v) glycerol gradients in the presence of molecular weight markers. After centrifugation, the fractions were analyzed by SDS-PAGE. The fractions at which the maximal amount of each protein appears are indicated by arrows.
(D) Comparison of the stimulatory effect of B35SSB/untagged B35SSB addition in M13 ssDNA replication assay. Reactions were carried out using primed M13 circular ssDNA as template and the Bam35 DNA polymerase in the presence or absence of B35SSB or untagged B35SSB as indicated. After incubation at 37 ºC, the length of the synthesized DNA was analyzed by alkaline 0.7% agarose gel electrophoresis and autoradiography.

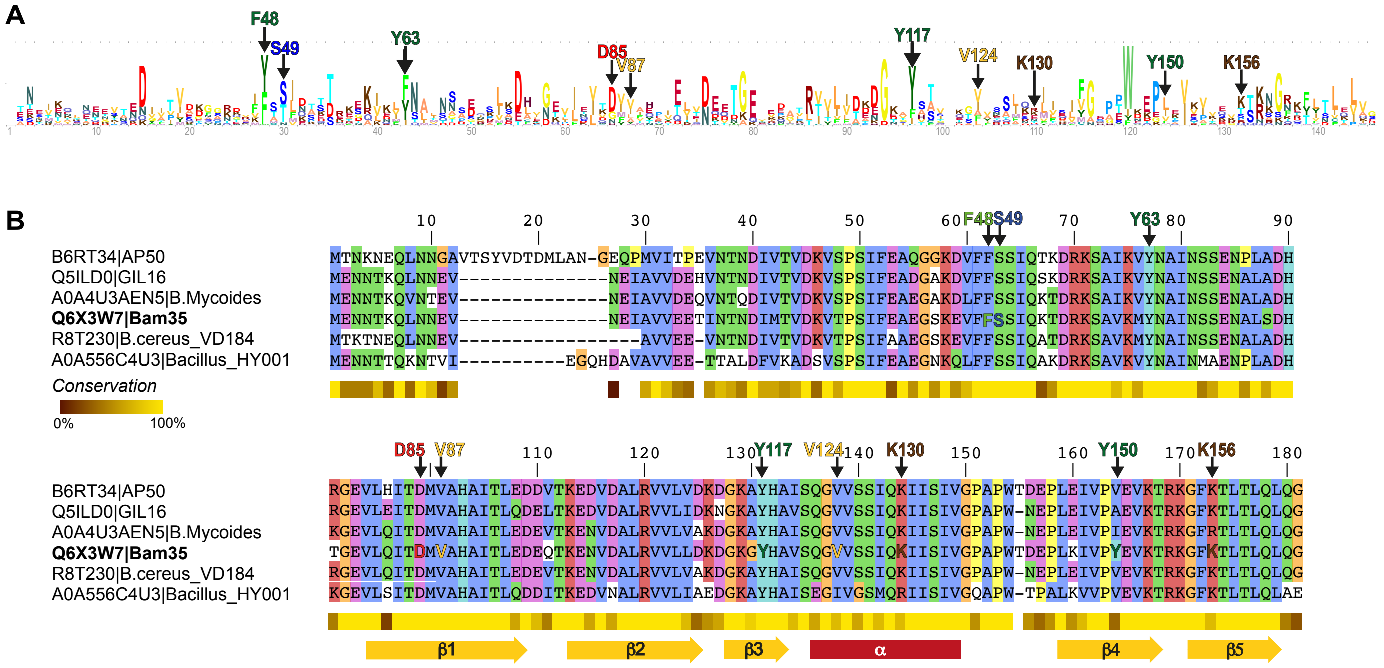


**Figure S3. Conserved residues in B35SSB-related proteins.**

(A) Logo of the conserved residues among Bam35-Φ29 SSBs. The Bam35-Φ29 SSBs logo was generated from the same alignment as shown in Figure 6. The frequency of each residue is represented by the height of that letter within a stack.

(B) Multiple sequence alignment of betatectiviral orthologs to B35SSB. The alignment was generated with MAFFT [44] and visualized with Jalview [45]. The level of conservation and predicted secondary structure of the C-terminal conserved fold of Bam35-Φ29 SSBs are shown. The B35SSB residues studied in this work are also highlighted.

<table>
<tr><th>No</th><th>Hit</th><th>Prob</th><th>E-value</th><th>P-value</th><th>Score</th><th>SS</th><th>Cols</th><th>Query HMM</th><th>Template HMM</th><th></th></tr>
<tr><td>1</td><td>PF17427.3 ; Phi29_Phage_SSB ;</td><td>100.0</td><td>5.9E-40</td><td>4.3E-45</td><td>253.9</td><td>18.0</td><td>124</td><td>42-166</td><td>2-125</td><td>(125)</td></tr>
<tr><td>2</td><td>ECOD_000118946_e1ffyA4 | 375.1</td><td>46.7</td><td>1.9E+02</td><td>0.0014</td><td>21.0</td><td>4.8</td><td>40</td><td>104-145</td><td>4-43</td><td>(66)</td></tr>
<tr><td>3</td><td>ECOD_001389154_e4u48A3 | 11.1.</td><td>33.5</td><td>2.3E+02</td><td>0.0017</td><td>20.4</td><td>3.7</td><td>30</td><td>91-120</td><td>15-44</td><td>(88)</td></tr>
<tr><td>4</td><td>4UAV_A CbbY; haloacid dehaloge</td><td>32.5</td><td>2.1E+02</td><td>0.0015</td><td>22.8</td><td>3.6</td><td>38</td><td>105-142</td><td>3-40</td><td>(246)</td></tr>
<tr><td>5</td><td>ECOD_001121967_e4m8oA1 | 2.1.1</td><td>32.3</td><td>3.4E+02</td><td>0.0025</td><td>23.9</td><td>4.9</td><td>33</td><td>88-120</td><td>52-102</td><td>(140)</td></tr>
<tr><td>6</td><td>PF15911.6 ; WD40_3 ; WD domain</td><td>31.2</td><td>1.3E+02</td><td>0.00098</td><td>21.6</td><td>2.2</td><td>14</td><td>104-117</td><td>20-33</td><td>(58)</td></tr>
<tr><td>7</td><td>d4uava_ c.108.1.0 (A:) automat</td><td>29.6</td><td>3.1E+02</td><td>0.0023</td><td>22.3</td><td>4.2</td><td>38</td><td>105-142</td><td>3-40</td><td>(246)</td></tr>
<tr><td>8</td><td>PF14433.7 ; SUKH-3 ; SUKH-3 im</td><td>28.6</td><td>1.9E+02</td><td>0.0014</td><td>23.7</td><td>3.0</td><td>35</td><td>99-133</td><td>103-144</td><td>(145)</td></tr>
</table>

**Figure S4. Results of a HHpred search using B35SSB as a query.**
B35SSB profile was generated by running eight iterations of PSI-BLAST against nr70 database.

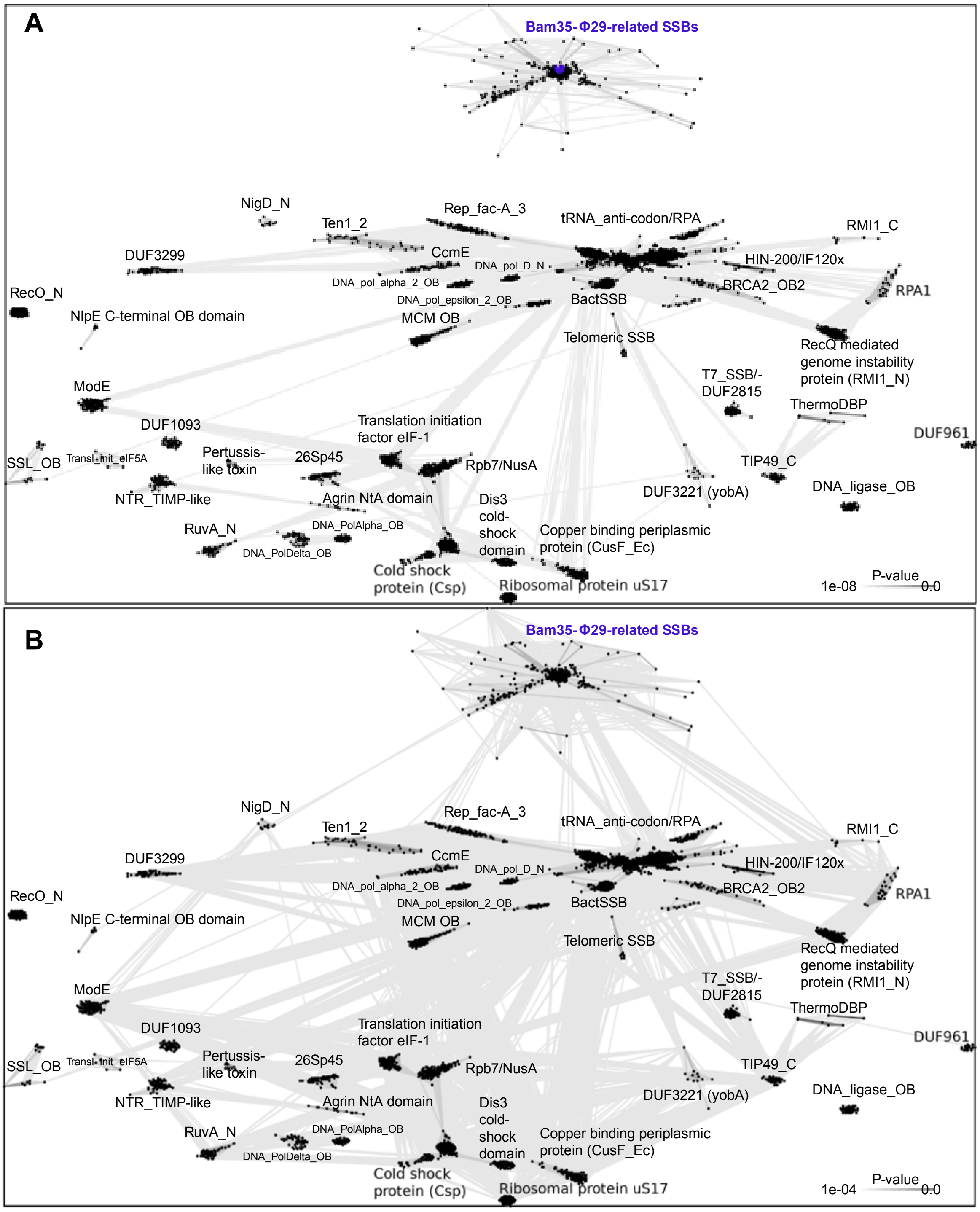


**Figure S5. All-to-all comparison of OB-fold-containing SSBs and Bam35-Φ29 SSBs.**

The dataset containing previously described OB-fold containing SSBs and Bam35-Φ29 SSBs was clustered using CLANS. Lines connect sequences with P-value ≤ $10^{-8}$ (A) or ≤ $10^{-4}$ (B), respectively. The obtained clusters are labeled. See Methods for details

```
# Job: B35SSB_cmRobetta
# Query: s001A
# No:  Chain   Z    rmsd lali nres  %id PDB  Description
   1:  4gnx-C  3.2  6.3   73   433    5   MOLECULE: PUTATIVE UNCHARACTERIZED PROTEIN;
   2:  5oun-A  3.2  2.5   64   107    9   MOLECULE: RUVB-LIKE PROTEIN 2;
   3:  3rd4-B  3.1  2.7   68    82   12   MOLECULE: UNCHARACTERIZED PROTEIN;
   4:  6fhs-C  3.1  4.0   66   459   17   MOLECULE: RUVB-LIKE HELICASE;
   5:  2k50-A  3.1  2.7   68   110    6   MOLECULE: REPLICATION FACTOR A RELATED PROTEIN;
   6:  4dni-A  3.1  4.0   66   257    9   MOLECULE: FUSION PROTEIN OF RNA-EDITING COMPLEX PROTEINS MP
   7:  1qvc-A  3.0  4.7   66   145    3   MOLECULE: SINGLE STRANDED DNA BINDING PROTEIN MONOMER;
```

```
# Job: B35SSB_trRosseta
# Query: s001A
# No:  Chain   Z    rmsd lali nres  %id PDB  Description
   1:  7bqk-A  3.3  3.3   51   459    6   MOLECULE: METHYLTRANSF_2 DOMAIN-CONTAINING PROTEIN;
   2:  4gxb-A  3.2  7.7   70   263    7   MOLECULE: SORTING NEXIN-17;
   3:  6f8e-A  3.2  2.9   67   121   12   MOLECULE: PLECKSTRIN HOMOLOGY DOMAIN;
   4:  1sc7-A  3.2  3.6   89   567    6   MOLECULE: 5'-D(*AP*AP*AP*AP*AP*GP*AP*CP*TP*T)-3';
   5:  4imh-A  3.2  4.5   89   347   10   MOLECULE: HEMIN DEGRADING FACTOR;
   6:  6tnt-A  3.0  5.5   69  1583    4   MOLECULE: ANAPHASE-PROMOTING COMPLEX SUBUNIT 1;
```

```
# Job: B35SSB_RaptorX
# Query: s001A
# No:  Chain   Z    rmsd lali nres  %id PDB  Description
   1:  1gpc-A  3.5  3.0   95   218   12   MOLECULE: PROTEIN (CORE GP32);
   2:  2ffg-A  3.4  3.8   65    80    8   MOLECULE: YKUJ;
   3:  2k50-A  3.2  2.7   68   110    6   MOLECULE: REPLICATION FACTOR A RELATED PROTEIN;
   4:  6fhs-C  3.2  4.7   70   459   14   MOLECULE: RUVB-LIKE HELICASE;
   5:  4lzi-A  3.1  7.0   69   277    6   MOLECULE: MULTICYSTATIN;
   6:  6wjv-A  3.1  3.5   79  1951   11   MOLECULE: DNA POLYMERASE EPSILON CATALYTIC SUBUNIT A;
   7:  6cqm-D  3.1  2.3   61    99    8   MOLECULE: SINGLE-STRANDED DNA-BINDING PROTEIN RIM1, MITOCHO
   8:  5da9-A  3.1  5.9   67   433   10   MOLECULE: PUTATIVE UNCHARACTERIZED PROTEIN,PUTATIVE UNCHARA
   9:  5jrk-A  3.0  5.0   88   695    7   MOLECULE: DIPEPTIDYL AMINOPEPTIDASES/ACYLAMINOACYL-PEPTIDAS
  10:  4gnx-C  3.0  5.9   79   433    6   MOLECULE: PUTATIVE UNCHARACTERIZED PROTEIN;
  11:  1wnh-A  3.0  5.7   67   220    9   MOLECULE: LATEXIN;
  12:  6s85-D  3.0  3.7   64   375    6   MOLECULE: NUCLEASE SBCCD SUBUNIT C;
  13:  4gxb-A  3.0  5.5   73   263    7   MOLECULE: SORTING NEXIN-17;
```

**Figure S6. Structural comparisons of B35SSB models and PDB database.**
Dali hits with Z-score above 3 are shown.

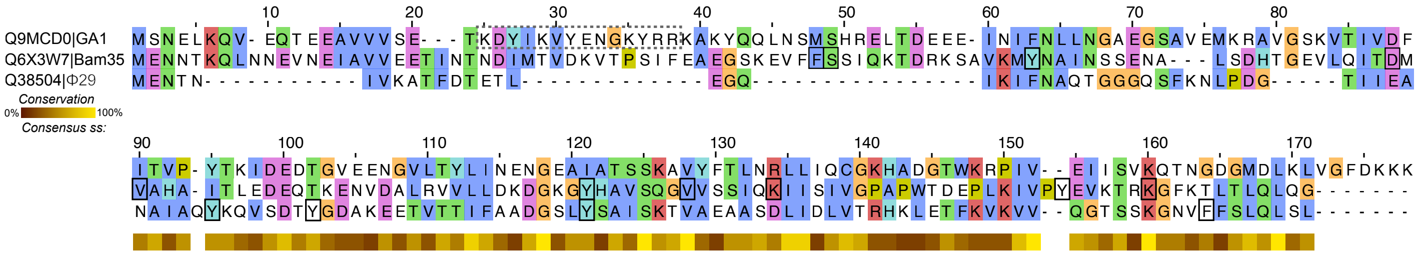


**Figure S7. Multiple sequence alignment of the Φ29, Bam35, and GA-1 viral SSBs.** MSA was generated with MAFFT [44] using E-INS-I method and visualized with Jalview [45]. The level of conservation is indicated below. B35SSB residues studied in this work, Φ29 characterized residues and GA-1 SSB N-terminal region essential for oligomerization are indicated. [38,66,74]. See text for details.

**Table S1. Oligonucleotides used in this work.**

Name, sequence and application are indicated. Sequences that correspond to restriction sites are highlighted by lowercase letter. The position of nuclease resistant phophorothioate bonds are indicated by an *.

| Name | Sequence (5'→ 3') | Application |
|---|---|---|
| B35SSB_FW_KpnI | GGCCggtaccAGATGGAAAACAATACTAAACAATTAAATAATGAGGTT | Generation of pET52b::B35SSB vector |
| B35SSB_RV_BamHI | CCCCggatccATTGCCTTGTAGTTGAAGTGTTAATGTTTTGAA | Generation of pET52b::B35SSB vector |
| B35SSB_D85A_FW | GTGAAGTTTTGCAAATTACGGCCATGGTGGCTCATGCTATTAC | B35SSB gene site-directed mutagenesis |
| B35SSB_D85A_RV | GTAATAGCATGAGCCACCATGGCCGTAATTTGCAAAACTTCAC | B35SSB gene site-directed mutagenesis |
| B35SSB_F48A_FW | GAAGGTTCAAAAGAAGTATTCGCCTCATCTATTCAAAAAACAGAC | B35SSB gene site-directed mutagenesis |
| B35SSB_F48A_RV | GTCTGTTTTTTGAATAGATGAGGCGAATACTTCTTTTGAACCTTC | B35SSB gene site-directed mutagenesis |
| B35SSB_K130A_FW | GGTGTTGTATCGTCTATTCAAGCGATTATCAGCATTGTTGGAC | B35SSB gene site-directed mutagenesis |
| B35SSB_K130A_RV | GTCCAACAATGCTGATAATCGCTTGAATAGACGATACAACACC | B35SSB gene site-directed mutagenesis |
| B35SSB_K156A_FW | CCTTATGAAGTAAAAACGCGTGCAGGTTTCAAAACATTAACAC | B35SSB gene site-directed mutagenesis |
| B35SSB_K156A_RV | GTGTTAATGTTTTGAAACCTGCACGCGTTTTTACTTCATAAGG | B35SSB gene site-directed mutagenesis |
| B35SSB_S49A_FW | GGTTCAAAAGAAGTATTCTTCGCATCTATTCAAAAAACAGACCG | B35SSB gene site-directed mutagenesis |
| B35SSB_S49A_RV | CGGTCTGTTTTTTGAATAGATGCGAAGAATACTTCTTTTGAACC | B35SSB gene site-directed mutagenesis |
| B35SSB_V124A_FW | CACGCCGTTTCACAAGGTGCTGTATCGTCTATTCAAAAG | B35SSB gene site-directed mutagenesis |
| B35SSB_V124_RV | CTTTTGAATAGACGATACAGCACCTTGTGAAACGGCGTG | B35SSB gene site-directed mutagenesis |
| B35SSB_V87A_FW | CAAATTACGGACATGGCGGCTCATGCTATTAC | B35SSB gene site-directed mutagenesis |
| B35SSB_V87A_RV | GTAATAGCATGAGCCGCCATGTCCGTAATTTG | B35SSB gene site-directed mutagenesis |
| B35SSB_Y117A_FW | GGACAAGGATGGAAAAGGTGCTCACGCCGTTTCACAAGG | B35SSB gene site-directed mutagenesis |
| B35SSB_Y117A_RV | CCTTGTGAAACGGCGTGAGCACCTTTTCCATCCTTGTCC | B35SSB gene site-directed mutagenesis |
| B35SSB_Y117F_FW | CAAGGATGGAAAAGGTTTTCACGCCGTTTCACAAG | B35SSB gene site-directed mutagenesis |
| B35SSB_Y117F_RV | CTTGTGAAACGGCGTGAAAACCTTTTCCATCCTTG | B35SSB gene site-directed mutagenesis |
| B35SSB_Y150A_FW | GAACCACTTAAAATTGTACCTGCTGAAGTAAAAACGCGTAAAGG | B35SSB gene site-directed mutagenesis |
| B35SSB_Y150A_RV | CCTTTACGCGTTTTTACTTCAGCAGGTACAATTTTAAGTGGTTC | B35SSB gene site-directed mutagenesis |
| B35SSB_Y150F_FW | CCACTTAAAATTGTACCTTTTGAAGTAAAAACGCGTAAAG | B35SSB gene site-directed mutagenesis |
| B35SSB_Y150F_RV | CTTTACGCGTTTTTACTTCAAAAGGTACAATTTTAAGTGG | B35SSB gene site-directed mutagenesis |
| B35SSB_Y63A_FW | CCGTAAATCAGCAGTTAAAATGGCCAATGCTATTAACTCAAGTG | B35SSB gene site-directed mutagenesis |
| B35SSB_Y63A_RV | CACTTGAGTTAATAGCATTGGCCATTTTAACTGCTGATTTACGG | B35SSB gene site-directed mutagenesis |
| B35SSB_Y63F_FW | CCGTAAATCAGCAGTTAAAATGTTCAATGCTATTAACTCAAGTG | B35SSB gene site-directed mutagenesis |
| B35SSB_Y63F_RV | CACTTGAGTTAATAGCATTGAACATTTTAACTGCTGATTTACGG | B35SSB gene site-directed mutagenesis |
| T7_FW | TAATACGACTCACTATAGGGCGA | Sequencing of pET52b::B35SSB vector and variants |

| | | |
|---|---|---|
| 15-mer | GATCACAGTGAGTAC | DNA binding assays |
| 33-mer | ACTGGCCGTCGTTCTATTGTACTCACTGTGATC | DNA binding assays |
| 50-mer | GAAAATAAAAATAAACTGGACAATTAAGGATCCTGTACAAGTCGACGCGG | DNA binding assays |
| 50-mer_c | CCGCGTCGACTTGTACAGGATCCTTAATTGTCCAGTTTATTTTTATTTTC | DNA binding assays |
| 50-mer(2) | GATCACAGTGAGTACAATAGAACGACGGCCAGTTGTCTCAATCTAACGGC | DNA binding assays |
| 80-mer | GGCCCATATGTACCCATACGACGTCCCAGACTACGCTGGACGGGGAGCGGGGCC AGTAAACTACT GACTGCGCAAGAGG | DNA binding assays |
| B35-ORI | TATTATGGTACCCCTACCAACCTATTAAA | DNA binding assays |
| B35-ORIc | TTTAATAGGTTGGTAGGGGTACCATAATA | DNA binding assays |
| RNA-50mer | GAAAAUAAAAAUAAACUGGACAAUUAAGGAUCCUGUACAAGUCGACGCGG | DNA binding assays |
| 15-mer | GATCACAGTGAGTAC | Cross-linking experiments |
| 17-mer | GATCACAGTGAGTACAA | Cross-linking experiments |
| 20-mer | GATCACAGTGAGTACAATAG | Cross-linking experiments |
| 21-mer | GATCACAGTGAGTACAATAGA | Cross-linking experiments |
| 22-mer | GATCACAGTGAGTACAATAGAA | Cross-linking experiments |
| 23-mer | GATCACAGTGAGTACAATAGAAC | Cross-linking experiments |
| 24-mer | GATCACAGTGAGTACAATAGAACG | Cross-linking experiments |
| 25-mer | GATCACAGTGAGTACAATAGAACGA | Cross-linking experiments |
| 30-mer | AACTGCCAAGAATAGTGTCAGTTCCAGACG | Cross-linking experiments |
| 40-mer | AGAGACAACTGGCCGTCGTTCTATTGTACTCACTGTGATC | Cross-linking experiments |
| 50-mer_b | GATCACAGTGAGTACAATAGAACGACGGCCAGTTGTCTCAATCTAACGGC | Cross-linking experiments |
| M13UP | GTAAAACGACGGCCAGT | M13 singly-primed replication |
| B35-ORIc_29mer | TTTAATAGGTTGGTAGGGGTACCATAA*T*A | Protein-primed replication initiation |
| B35-ORIc_60mer | TAAATGTTATTACATGTCAATAGGTAGATATTTTAATAGGTTGGTAGGGGTACCATAA* T*A | Protein-primed replication initiation |
